# Embryonic Signatures of Intergenerational Inheritance across Paternal Environments and Genetic Backgrounds

**DOI:** 10.1101/2024.11.25.624914

**Authors:** Mathilde Dura, Bobby Ranjan, Rossella Paribeni, Violetta Paribeni, Joana B. Serrano, Laura Villacorta, Vladimir Benes, Olga Boruc, Ana Boskovic, Jamie A. Hackett

## Abstract

The paternal preconception environment has been implicated as a modulator of phenotypic traits and disease risk in F_1_ offspring. However, the prevalence and mechanisms of such intergenerational epigenetic inheritance (IEI) in mammals remain poorly defined. Moreover, the interplay between paternal exposure, genetics, and age on emergent offspring features is unexplored. Here, we measure the quantitative impact of three paternal environments on early embryogenesis across genetic backgrounds. Using *in vitro* fertilisation (IVF) at scale, we capture batch-robust transcriptomic signatures of IEI with single-blastocyst resolution. Amongst these, paternal gut microbiota dysbiosis is linked with aberrant expression of lineage regulators in blastocysts, particularly affecting extra-embryonic tissues. Conversely, paternal low-protein high-sugar diet associates with subtle preimplantation developmental delay. We further identify gene expression variability as a paternally-induced F_1_ phenotype, and highlight confounding issues for IEI such as batch-effects and under-sampling. Finally, paternal genetic background and age exert a dominant influence over the inherited environmental signature. This study systematically characterises how paternal conditioning programmes subtle but detectable molecular responses in early embryos, and suggests guiding principles to dissect intergenerational phenomenology.

## Introduction

Inheritance of biological material entails the transmission of genetic and non-genetic information across generations. In mammals, mature sperm transfer genomic DNA sequence to oocytes during fertilisation, which serves as the fundamental basis of paternal inheritance (Watson & Crick, 1953). In addition, sperm also transmit ‘epigenetic’ information in the form of small RNA payloads, chromatin states, DNA methylation, endocrine signals and metabolites (Adrian-Kalchhauser *et al*, 2020; Fitz-James & Cavalli, 2022; Skvortsova *et al*, 2018). These non-genetic molecular factors have the potential to influence early embryonic genome regulation, and thus to affect development, and ultimately F_1_ phenotype. Moreover, epigenetic information in the germline is relatively dynamic compared to the DNA sequence, with the potential to change in response to extrinsic cues and/or environmental perturbations (Ciabrelli *et al*, 2017; Murphy *et al*, 2020; Rechavi *et al*, 2014; Torres-Garcia *et al*, 2020). As a consequence, an increasing number of mammalian studies report that fathers exposed to adverse pre-conception environments transmit phenotype(s) to their F_1_ offspring, referred to as intergenerational epigenetic inheritance (IEI). Such intergenerational epigenetic effects that traverse a single generation are emerging as important but overlooked contributors to phenotypic variation and disease susceptibility, yet remain poorly understood.

Multiple paternal environments have been reported to propagate offspring effects, including dietary perturbations (high-fat, low-protein, folate-deficiency), drug exposure (smoking, antidiabetics), gut dysbiosis (antibiotics) and other extrinsic stressors (trauma, endocrine disrupting chemicals) (Argaw-Denboba *et al*, 2024; Carone *et al*, 2010; Chen *et al*, 2016; Huypens *et al*, 2016; Lambrot *et al*, 2013; Lesch *et al*, 2019; Radford *et al*, 2014; Tomar *et al*, 2024; Vallaster *et al*, 2017). Amongst these, perturbations to the gut microbiome of prospective fathers have been shown to influence the birthweight and mortality rate of offspring (Argaw-Denboba *et al*., 2024; Masson *et al*, 2024). This effect occurs probabilistically and manifests as an increased risk of adverse F_1_ outcomes rather than a deterministic response. Mechanistically, dysbiotic fathers affect offspring by inducing *in utero* placental insufficiency, indicating the origin of emergent F1 phenotypes can be traced back to early embryogenesis. Indeed, prior studies found that offspring of fathers exposed to low-protein high-sugar (LPHS) diet exhibit altered expression of hepatic lipid-associated genes and metabolic dysfunction (Carone *et al*., 2010). This was linked with paternally-induced changes in MERVL expression as early as preimplantation development (Sharma *et al*, 2016). It is now crucial to obtain a fuller understanding of the presentation of molecular signatures in early embryos, towards dissecting the underlying mechanisms.

IEI does not occur *in silo*, and numerous factors can affect its propagation and/or manifestation. For instance, environmental exposures are interconnected, raising the question of whether each exposure produces a distinct outcome or whether they funnel into a limited number of generic F_1_ responses. For example, antibiotics and LPHS administration both impact gut microbiome composition (Ross *et al*, 2024), but also exert treatment-specific effects on fathers and offspring (Argaw-Denboba *et al*., 2024; Carone *et al*., 2010). Beyond this bandwidth of potential F_1_ responses, epigenetic phenomena are tightly coupled with genetics, implying that IEI might be contingent on the underlying DNA sequence (Cavalli & Heard, 2019). Consistently, phenotypic traits in individuals stem from interactions between their genetics and environment (GxE), and this multi-factorial relationship likely also modifies the signature and prevalence of IEI, yet is poorly understood. Indeed, there is currently limited understanding of precisely which epigenetic factors are responsible for transmitting IEI phenotypes and their molecular manifestation in embryos (Santilli & Boskovic, 2023). One hypothesis is that subtle perturbations introduced by exposed sperm into fertilised eggs snowball through successive stages of development, ultimately reaching a threshold that triggers physiological abnormalities. In order to disentangle these complex relationships, a high-throughput quantitative readout is necessary to capture the earliest molecular signatures of mammalian IEI across diverse GxE contexts.

To this end, we designed a large-scale multi-paradigm study profiling intergenerational responses in preimplantation embryos across paternal exposures, ages and genetic backgrounds. We used *in vitro* fertilisation (IVF) coupled with high-throughput transcriptomic profiling of preimplantation blastocysts to ask the following questions: (i) Can we capture consistent and reproducible molecular signatures of IEI, (ii) Are these signatures common or specific to paternal exposure, and do they imply mechanisms/outcomes, (iii) How do paternal genetic background and age influence these signatures, and (iv) Are F_1_ signatures detectable upstream in the paternal reproductive system?

## Results

### An IVF-based multi-paradigm model of paternal effects

We first sought to establish a tractable experimental model that can capture the earliest molecular signatures of paternal environments on progeny. We designed a multi-paradigm IVF-strategy with single-embryo transcriptomics, using sperm from males exposed to distinct environmental factors (Fig 1A). IVF enables replicates of independent embryos to be generated at scale within a highly controlled context, across multiple parallel batches, thus increasing power whilst controlling for experimental variables. Specifically, our IVF strategy rules out confounders from natural matings such as maternal resource allocation based on judgment of male quality, indirect effects of paternal perturbation on females (*e*.*g*. via coprophagy), and/or litter-size effects (Bohacek & Mansuy, 2017). As paternal environment perturbations, we chose to focus on factors previously linked with inducing intergenerational postnatal phenotype(s), by exposing prospective fathers to: (i) non-absorbable antibiotics (nABX) to induce gut microbiome dysbiosis, (ii) low-protein high-sugar (LPHS) isocaloric western-style diet, and (iii) control standard diet (CON). Male FVB mice were thus exposed to nABX, LPHS or control diet/water *ad libitum* from weaning for 6-7 weeks (**Fig 1A**).

**Figure 1.**
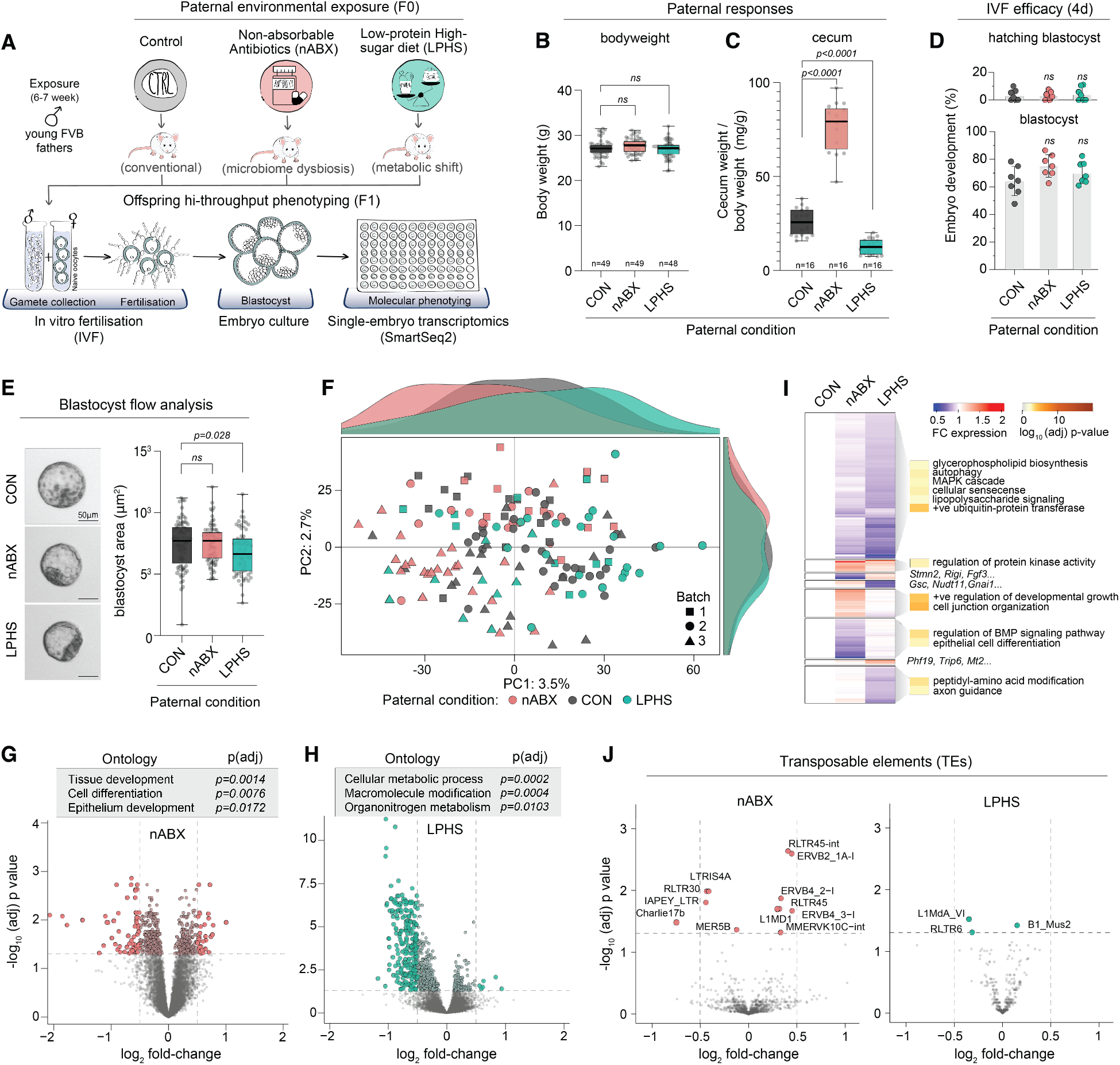
Offspring signatures of paternal environmental exposure. (A) Schematic of the experimental design. Young FVB males are exposed to non-absorbable antibiotics (nABX) or low-protein high-sugar (LPHS) diet for 6-7 weeks. Purified sperm are then collected for parallel *in vitro* fertilisation (IVF), with the resulting blastocysts undergoing single-embryo transcriptomics using smart-Seq2. (B-C) Box plots showing paternal F0 physiological responses to indicated treatment for (B) bodyweight and (C) cecum weight relative to bodyweight. Each datapoint represents a biological replicate (individual mouse), bars represent the median, and whiskers the minimum-maximum range. (D) Dot plot showing efficiency of development to blastocyst (bottom) and hatching blastocyst (top) depending on paternal sperm donor condition. (E) Representative images of blastocysts from the three paternal conditions (left), and box plot showing blastocyst size (area) by paternal condition. (F) Principal component analysis (PCA) of blastocyst global transcriptome (all expressed genes) coloured according to paternal treatment. Each data point indicates a single blastocyst and histograms represent relative density of blastocysts along PC1 (top) and PC2 (right). (G-H) Volcano plots showing differential gene expression in (G) nABX and (H) LPHS derived blastocysts. Grey boxes show gene ontology (GO) terms significantly enriched in the respective differentially expressed gene sets. (I) Heatmap showing average expression fold-change of DE genes clustered according to F1 response pattern. Shown right selected GO terms significantly enriched in each DE gene cluster. (J) Volcano plots depicting differential expression of transposable elements in paternal nABX (left) or LPHS (right) derived blastocysts. (B-E) P-values were computed using a two-tailed unpaired t-test [ns: not significant].

We first scored the direct impact of environmental factors on male physiology. Neither nABX nor LPHS impacted survival, overt phenotype(s) or body weight after six weeks. We did however note a reciprocal trend towards increased (nABX) and decreased (LPHS) body mass (**Fig 1B, Fig S1A-C**). Moreover, we found major responses within the gastrointestinal tract. Specifically, gut dysbiosis induced by nABX was associated with significantly increased cecum size and weight, whereas in contrast, LPHS diet led to a reduction in caecum weight (**Fig 1C, Fig S1D-E**). Such effects imply the two environmental paradigms have at least partially reciprocal impacts on systemic physiology in prospective fathers.

We next asked how paternal exposures impact fertilisation efficacy. High quality sperm from treated and control males were isolated after swim-up, and IVF was performed from a randomised common pool of strain-matched oocytes for all sperm donors. All IVFs were performed in parallel on the same morning to minimise intra-batch variability. The resulting zygotes were cultured *ex vivo*, with no differences observed in the rate of blastocyst development or hatching across the paternal conditions (**Fig 1D**). While blastocysts from all paternal treatments appeared morphologically indistinguishable, using quantitative whole-embryo flow analysis, we noted a small but significant decrease in the size of blastocysts derived from LPHS-exposed males, pointing to a potential developmental delay (**Fig 1E; Fig S1F-H**). In summary, we established a paternal multi-paradigm exposure model coupled with controlled and scalable *ex vivo* readouts of early F_1_ effects. We observe direct physiological responses in exposed males, but no gross impact on fertility or rates of F_1_ embryonic development.

### Intergenerational molecular signatures of paternal environments

To investigate whether the impacts of paternal environment could manifest as an altered molecular profile in early embryos, we undertook single-embryo transcriptome profiling. To capture the earliest effects we assayed blastocysts, which represent the first developmental stage after embryonic and extra-embryonic lineages are established, can be cultured at scale, and can be accurately staged. These considerations minimise variation related to rapid developmental transitions during cleavage divisions, whilst enabling putative effects on early lineages to manifest, thus making blastocysts ideal to capture robust and consistent transcriptomic signatures of IEI. Individual blastocysts derived from the three paternal conditions were sequenced with SMART-seq2, yielding high-quality, high-coverage transcriptomes from 157 embryos, each with 5-20 million reads (**Fig S1I-K**). The male-female ratio was broadly even across conditions (**Fig S1L**).

Global analysis of all expressed genes by principal component analysis (PCA) revealed blastocysts did not cluster by paternal treatment, consistent with no overt developmental F_1_ phenotype. However, we did observe a tendency of nABX-and LPHS-derived blastocysts to separate away from control embryos, with the respective exposures moving in opposite directions along the first principal component (PC1) of transcriptome variation (**Fig 1F**). Indeed, the density of blastocyst projections along PC1 exhibited a trimodal distribution that reflected paternal nABX, LPHS or control status at conception (**Fig 1F**). These data imply that while there is no deterministic global response to paternal condition, a probabilistic tendency towards specific molecular effects in F_1_ embryos can be observed.

To further investigate this finding, we identified significant differentially expressed genes (DEG) from each paternal condition, leveraging high replicate numbers across multiple batches to ensure robustness. This revealed that embryos derived from nABX or LPHS fathers exhibit a high-confidence (hc) DEG profile (adjusted p<0.05, log_2_ FC>0.5) (**Fig 1G-H; Table S1**). Specifically, we found 97 hcDEG in nABX-fathered blastocysts, which were enriched for functional roles in cell differentiation and developmental pathways, implying paternal dysbiosis may be linked with an F_1_ lineage effect (**Fig 1G**). Amongst the most significantly effected genes were key cell fate regulators *Dkk1, Gata4* and *Tbx20*. In contrast, hcDEG in LPHS-derived blastocysts (*n=244*) were enriched for cellular metabolic processes and were mostly downregulated (**Fig 1H**). This included downregulation of many ribosome-associated genes such as *Rps2, Fbl* and *Utp18*, which is rate-limiting for growth (Buszczak *et al*, 2014). Considered together with significantly smaller blastocyst size (**Fig 1E**), these data are consistent with a developmental growth delay of embryos derived from LPHS fathers. Overall, our results indicate we can capture a robust molecular signature of paternal conditioning in early embryos.

Notably, there was little overlap between the two sets of hcDEG (14 genes), suggesting that distinct paternal perturbations induce distinct F_1_ effects at the blastocyst stage (**Fig S1M**). This is in line with the reciprocal physiological response of fathers to each treatment, the opposing trend of global blastocyst transcriptomes along PC1 (**Fig 1C, F**), and the divergent gene sets enriched across the conditions (**Fig S1N-O**). To further understand the specificity of F_1_ responses, we clustered all DEGs by their fold-change across conditions, obtaining eight clusters (Fig 1I). Genes specifically downregulated in nABX-derived blastocysts (cluster 6) were highly enriched for bone morphogenetic protein (BMP) signalling, which plays a key role in extra-embryonic lineage allocation (Graham *et al*, 2014), while nABX-specific upregulated genes (cluster 5) were linked with cell junction proteins and epithelialisation. The genes preferentially downregulated in LPHS-derived blastocysts (cluster 1) were enriched for senescence, ubiquitin activity and the MAPK cascade, which has been implicated in implantation failure (Natale *et al*, 2004). In this case, genes determined to be preferentially LPHS-specific also trend towards downregulation in the nABX paradigm, hinting at a partially shared F_1_ response of different magnitudes (**Fig 1I**). Overall, the data suggest that while some of the intergenerational responses may reflect a general effect of adverse paternal conditioning, specific paternal exposures also trigger specific pathway responses as early as the blastocyst stage.

Finally, we investigated the paternal impact on transposable element (TE) expression in offspring. TEs have been implicated as both markers and functional regulators of successive stages of preimplantation development, and have also been previously linked with epigenetic inheritance (Modzelewski *et al*, 2021; Morgan *et al*, 1999; Todd *et al*, 2019). We observed only limited significant TE responses in LPHS-derived blastocysts (**Fig 1J; Table S2**). However, we note significant changes in TE expression in nABX-derived blastocysts, primarily in long terminal repeat (LTR) retrotransposons, including upregulation RLTR45 and MMERVK10C and downregulation of IAPey (**Fig 1J; Table S2**). These TEs are dynamically expressed during embryonic progression, and have been shown to be amongst the most epigenetically responsive TEs (Hackett *et al*, 2017; Reichmann *et al*, 2012), pointing towards an aberrant epigenome and/or differences in development. Taken together, we were able to identify significant albeit subtle F_1_ signatures specific to each paternal exposure, reflecting both protein-coding genes and transposable element responses.

### The regulatory basis of intergenerational signatures

To investigate potential mechanisms underlying F_1_ gene expression changes, we assessed the upstream regulators of DEGs by searching for transcription factors (TFs) that bind their proximal promoters. We found TFs that confer lineage identity, such as NANOG and ZIC3, to be highly enriched at promoters of nABX-specific DEGs (**Fig 2A; Fig S2A**). Moreover, the downstream effectors of BMP signalling, SMAD1, SMAD3 and SMAD4, were also amongst the most significant binders of DEGs, consistent with enrichment for BMP signalling pathways in nABX blastocysts. In contrast, DEGs from LPHS blastocysts were associated with MYC and TRIM28 promoter-proximal binding (Fig 2A), which are linked with cell fitness/proliferation and targeting H3K9me3, respectively (Claveria *et al*, 2013; Rowe *et al*, 2013; Scognamiglio *et al*, 2016). To understand whether F_1_ DEGs also exhibit a chromatin signature that could give us insight into their regulation, we leveraged published ChIP-Seq datasets from the ICM of blastocysts (Liu *et al*, 2016; Wang *et al*, 2018; Xu *et al*, 2019). We observed an enrichment of H3K9me3 preferentially in the promoters of LPHS-DEGs, consistent with those being TRIM28 targets (**Fig 2B**). Curiously, such H3K9me3 often co-occurred with H3K4me3, with this chromatin signature being absent in the promoters of nABX-specific DEG (**Fig 2B-C**). The cumulative data imply that paternal LPHS diet may influence F_1_ regulatory pathways related to growth and proliferation, whereas dysbiotic nABX-exposed fathers could affect offspring pathways involved in cell-lineage allocation.

**Figure 2.**
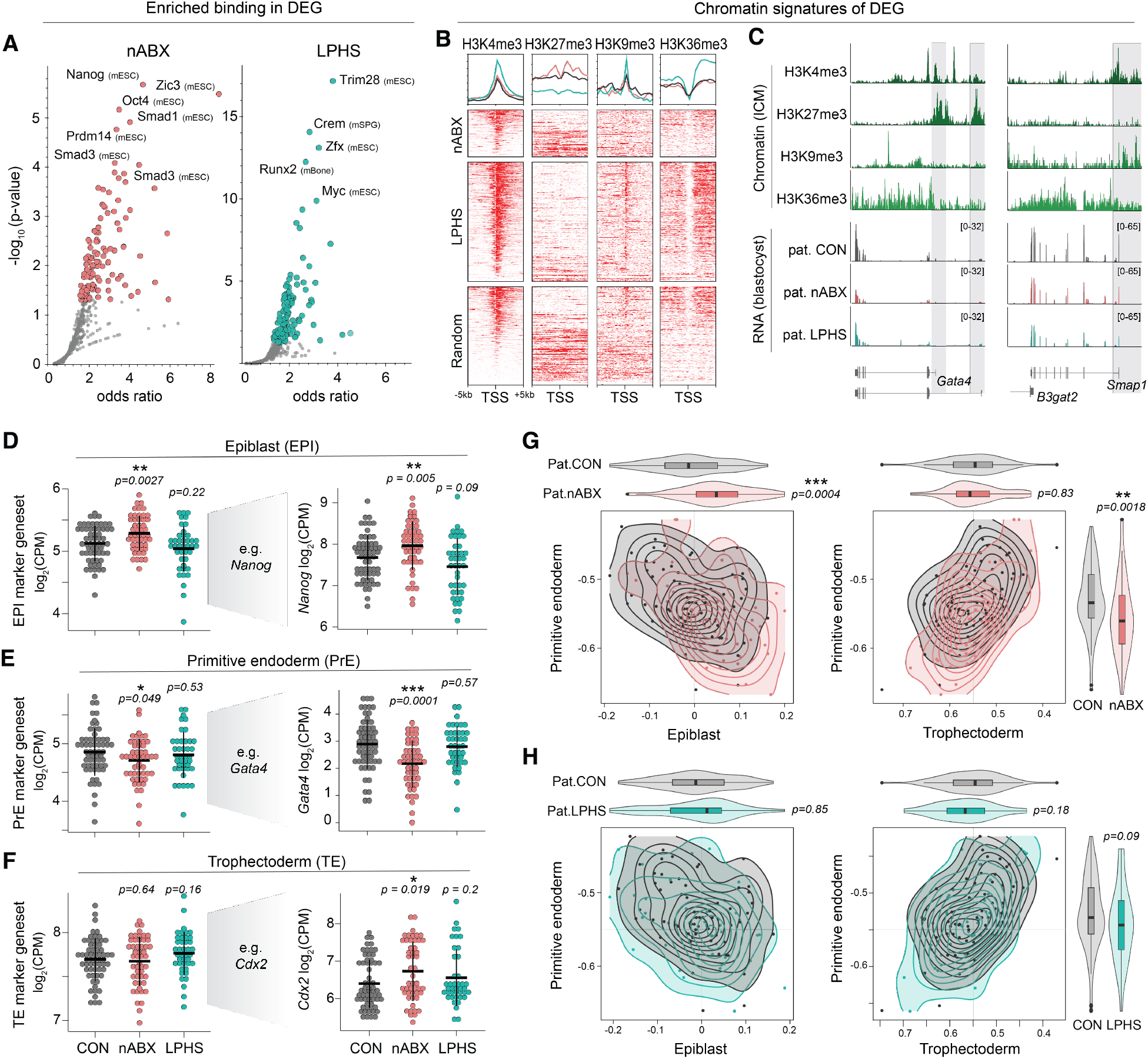
The regulatory basis of intergenerational transcriptomic signatures. (A) Scatter plot showing enrichment for binding of specific transcription factors within DEG set promoters from paternal nABX- (left) or LPHS- (right) blastocysts. (B) Heatmaps and summary profiles of H3K4me3, H3K27me3, H3K9me3 and H3K36me3 enrichment around the TSS of DEG sets in wildtype blastocysts, sorted by H3K4me3 enrichment. Random genes are shown for comparison. (C) Representative genome tracks of DE genes showing chromatin marks in wildtype blastocysts and gene expression responses across the paternal treatments. (D-F) Dot plots depicting expression of lineage markers in embryos derived from each paternal condition. Shown left is mean expression of collective marker genes for each lineage: epiblast (top), primitive endoderm (middle) and trophectoderm (bottom). Markers used for the respective geneset (*see* Fig S2D). Shown right are representative genes for each lineage. Each datapoint indicates log2 CPM expression of a geneset/gene in a single blastocyst. *P*-values computed using two-tailed Wilcoxon test. (G-H) Contour plots and violin plots of correlation coefficients of blastocyst transcriptomic profiles from indicated paternal sperm donor with the reference single-cell lineage, comparing (G) nABX versus CON, and (H) LPHS versus CON. Correlation coefficients of primitive endoderm and epiblast (left) and primitive endoderm and trophectoderm (right) are displayed.

The appropriate formation and allocation of the three primary lineages in the developing blastocyst – epiblast (EPI), primitive endoderm (PrE) and trophectoderm (TrE) -is crucial to ensure implantation success. Indeed, recent studies have demonstrated that aberrant primitive endoderm is the key predictor of poor outcomes for human embryos (Chousal *et al*, 2024). To investigate whether paternal condition could trigger lineage-related defects, we grouped key markers for each lineage into genesets and measured their collective expression change as proxies for lineage allocation (see Methods). Blastocysts sired by nABX-treated males exhibited significant downregulation of primitive endoderm markers (*p=0*.*049*) and concomitant upregulation of epiblast genesets (p=*0*.*0027*) (Fig 2D-E). This was exemplified by *Nanog* and *Prdm14* for EPI, and *Gata4*, as well as its upstream regulator *Lrrc34* for PrE (Fig 2D-F, Fig S2A). We also observed that critical regulators for trophectoderm such as *Cdx2* were significantly mis-regulated in nABX embryos (Fig 2F). In contrast, we detected no lineage bias in blastocysts from LPHS sperm donors, which were concurrently derived from the same oocyte pool.

To further investigate the potential lineage bias in nABX-derived embryos, we constructed reference signatures of the three lineages using published single-cell blastocyst data (Deng *et al*, 2014) (**Fig S2B-D**), and projected our blastocyst profiles onto this reference. We observed an increased epiblast signature and decreased primitive endoderm signature specifically in paternal nABX-derived blastocysts (**Fig 2G-H**). Cumulatively, using multiple lines of transcriptomic enquiry, we observed lineage-specific expression changes in nABX-derived blastocysts, which manifest as decreased primitive endoderm marker expression coupled with increased epiblast marker activity. This could reflect a change in the proportion of cells allocated to each lineage and/or mis-regulation of lineage-genes themselves.

### Batch effects & design principles to capture intergenerational signatures

Early embryonic gene expression is inherently noisy, and capturing the effects of relatively subtle paternal influences is confounded by both technical and biological variation. In order to obtain a more robust picture of the intergenerational signatures, we profiled three independent batches of blastocysts across the three paternal environments. Each batch was designed such that all IVF and embryo culture stemming from different sperm donors occurs in parallel, using a shared oocyte pool, to minimise intra-batch confounders (**Fig 3A**). While prior analysis of integrated batches captured paternal-effect signatures (**Fig 1G-I**), we also found blastocysts preferentially separated according to batch, implicating batch as the strongest variable (**Fig 3B**). When not accounted for, batch effects can obscure *or* over-emphasise underlying biology.

**Figure 3.**
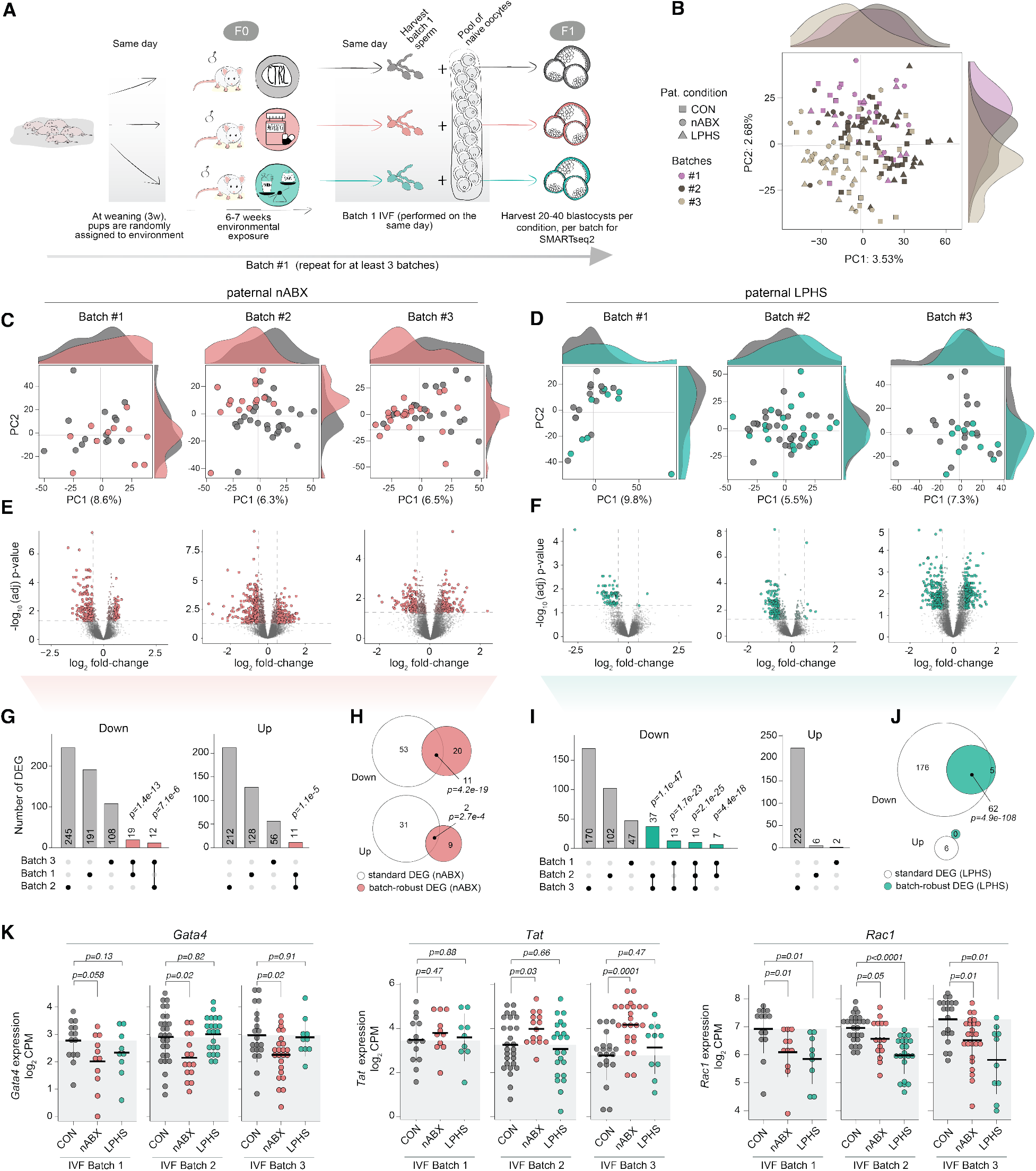
Overcoming batch effects to capture consistent IEI signatures. (A) Schematic depicting the organization of batches to obtain statistical power. IVFs from all sperm donors are performed on the same oocyte pool in parallel to generate embryos, which is repeated on multiple occasions (batches). (B) PCA on global transcriptomes coloured by batch. (C-D) PCA on global transcriptomes of F1 blastocysts separated by individual experimental batches, derived from (C) paternal nABX or (D) paternal LPHS exposures and relative to control fathers. Shaded histograms represent relative density of blastocysts along PC1 (top) and PC2 (right). (E-F) Volcano plots of DE genes from nABX or LPHS paternal treatment relative to controls in the same batch. (G) Upset plots comparing differential expressed gene (DEG) sets in each batch in the nABX model. (H) Venn diagrams showing overlap of upregulated (top) and downregulated (bottom) DE genes in the standard versus batch-robust analysis (broDEG). (I) Upset plots comparing differential expressed gene sets in each batch in the LPHS model. (J) Venn diagrams showing overlap of DE genes in the standard analysis versus batch-robust analysis. (G-J) P-values computed using Fischer’s exact test. (K) Dot plot of representative examples of DE genes that show consistent differential expression across batches. *P*-values indicate two tailed Wilcoxen test.

Our high replicate numbers allowed us to analyse each batch independently to mitigate confounding effects arising from inter-batch variation. PCA of each batch revealed a greater level of separation between F_1_ blastocysts from treated and control males than observed in the integrated analysis, particularly for paternal nABX treatment (**Fig 3C-D**). Moreover, independent batch analysis identified a marked number of hcDEG in blastocysts sired by nABX-or LPHS-exposed males relative to control (**Fig 3E-F, Fig S3A**). We used these to identify ‘batch robust’ DE genes (broDEG), comprising only genes that were DE in at least two batches (**Fig 3G-J1; Table S3**). The overlaps between such broDEGs and hcDEGs (**Fig 1G-I**) were modest but highly statistically significant (**Fig 3H-J**), indicating that the two strategies for identifying consistent transcriptomic changes can be used complementarily. Indeed, *Gata4* was systematically downregulated in nABX-derived blastocysts across all three batches. Moreover, *Lrrc34*, which regulates the expression of *Gata4*, along with *Nanog, Tat* and *Prdm14* were also consistently dysregulated across batches derived from dysbiotic fathers (**Fig 3K; Fig S3A-B**). Finally, developmental signalling genes like *Rac1* and *Mapk1* were reproducibly downregulated in LPHS samples (**Fig 3K**).

In summary, multiple paths of analysis led us towards the same conclusion: paternal nABX and LPHS treatments propagate information to the next generation that can be detected as gene expression changes in the F1 blastocysts. However, our analyses indicate that blastocyst expression profiles inherently possess a low signal-to-noise ratio (signature of paternal effect *vs* technical and batch-specific variation) for transmitted effects. This emphasises that paternal effects are subtle at the molecular level, and motivates the need for careful experimental design to guard against technical confounders or overinterpretation of intergenerational phenotypes.

### Interactions between paternal genetic background, age, & environment

Epigenetic phenomena are inextricable from genetics (Cavalli & Heard, 2019). Using inbred mouse strains provides an opportunity to assess, in a controlled manner, the role of <u>g</u>enetic background on the propagation of an environmentally-induced F_1_ phenotype (intergenerational GxE). To investigate this, we repeated the paternal exposure paradigms, IVF and blastocyst transcriptomic profiling using independent batches of C57BL/6 (B6) genetic background males as sperm donors and strain-matched oocytes. The direct physiological response of males was similar between the FVB and B6 backgrounds, with negligible impact on overt phenotypes and bodyweight, but opposite and significant differences in caecum weight (**Fig S4A-C**). Moreover, there was no significant difference across paternal conditions in developmental rate, hatching or male-female ratio of IVF-derived blastocysts (**Fig S4D-H**). We did however note that IVF using B6 sperm donors was markedly less efficient and more variable than FVB, irrespective of paternal conditioning, implying greater technical noise in the B6 background.

As observed in the FVB background, there was no global separation between transcriptomes of blastocysts derived from different paternal environments by PCA (**Fig 4A**). However, we also observed no significant differential expression across both paternal conditions, indicating the absence of a DEG signature. Genes reported to be DE in FVB did not show condition-specific expression directionality in the B6 background (**Fig 4B**). Similarly, differences in expression of lineage-associated markers did not reach significance across the conditions (**Fig 4C**). Using Gene Set Enrichment Analysis (GSEA), we observed that terms related to sodium ion transport, oxidative phosphorylation, and ribosomal activity, that were significantly enriched in the FVB background, showed little to no enrichment in B6 background (**Fig 4D**). In summary, the same environmental exposures in different genetic backgrounds lead to divergent intergenerational outcomes. Whilst this could be due to a range of biological or technical reasons (*see* Fig 5), it also highlights the significance of GxE interactions in intergenerational models, and implies that modifying either factor alters (or ablates) outcomes.

**Figure 4.**
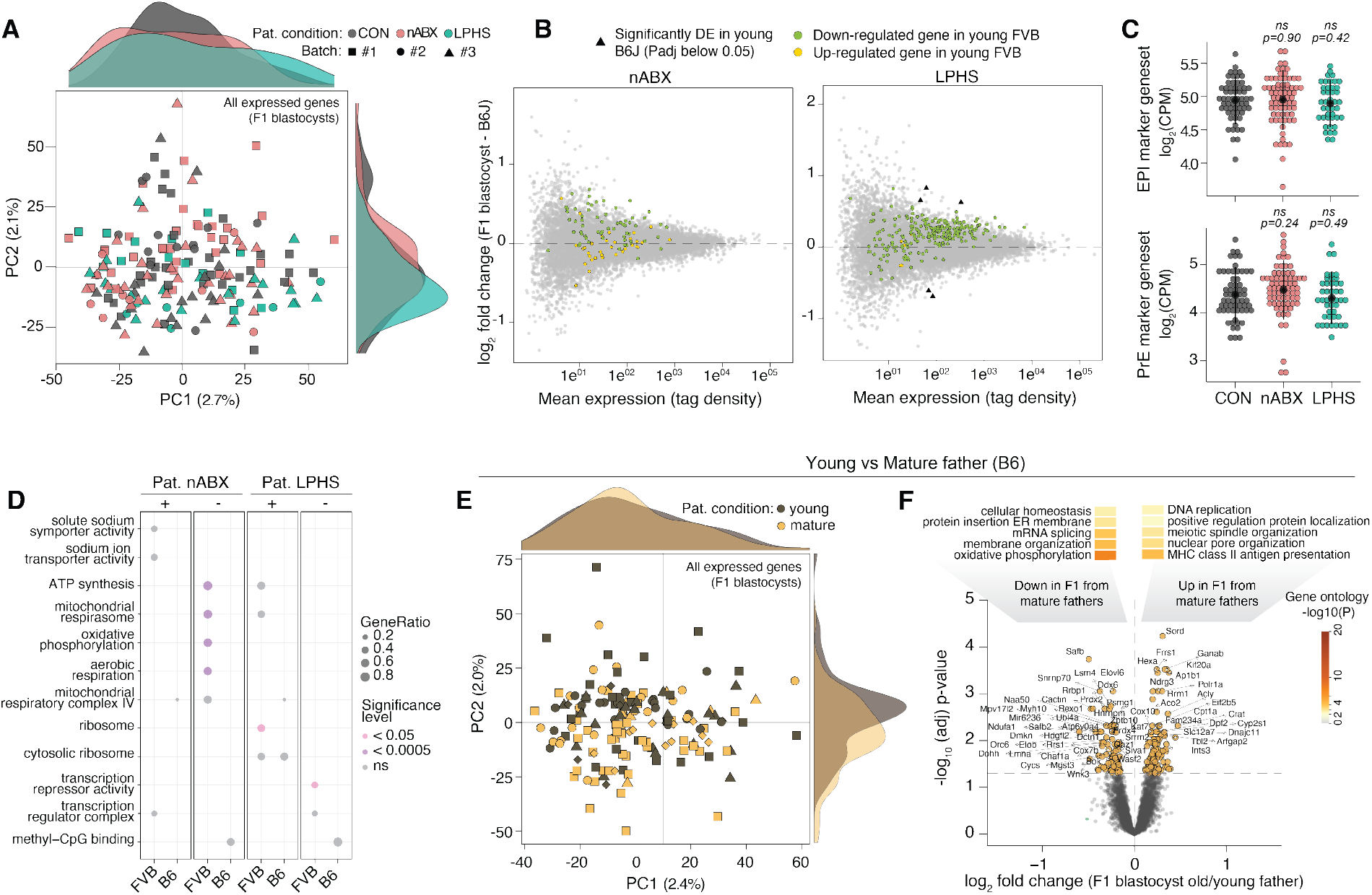
The impact of genetic background and age on intergenerational epigenetic inheritance. (A) PCA plot of blastocyst transcriptomes of C57BL/6 (B6) background across the three paternal treatments. (B) MA plot of (left) nABX and (right) LPHS blastocyst gene expression relative to B6 controls, showing relative expression of the IEI signature identified in the FVB genetic background. (C) Total expression (log2CPM) of selected epiblast (EPI) (top) and primitive endoderm (PrE) (bottom) markers. Markers used for the respective geneset (*see* Fig S2D). P-values were computed using a two-tailed Wilcoxon test. (D) Bubble plot showing gene set enrichment analysis (GSEA) comparing the paternal treatments across FVB and B6 backgrounds. (E) PCA on whole transcriptomes of blastocysts from young and mature B6 males. (F) Volcano plot showing differential expression between blastocysts derived from mature and young males. Shown above is a deconstructed heatmap with GO terms enriched in DE genes down-regulated (left) and up-regulated (right) in blastocysts from mature fathers. Thresholds for significance are set at adjusted p-value < 0.05 and abs(log2FoldChange) > 0.5.

**Figure 5.**
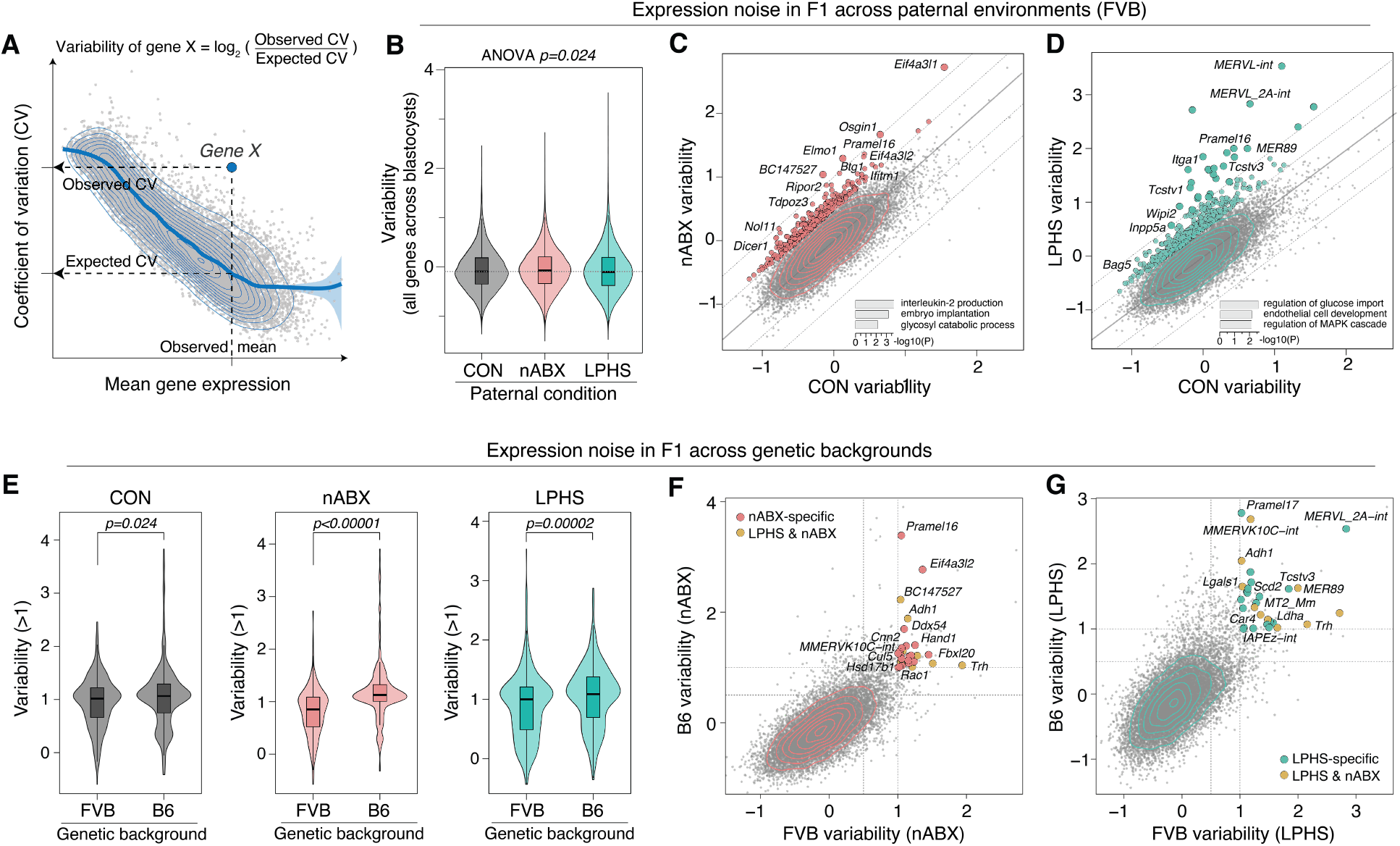
Intrinsic population variability as a readout of intergenerational epigenetic inheritance. (A) Scatter schematic depicting the calculation of variability of a gene “X” using coefficient of variation (CV) versus mean expression. (B) Violin and box plots comparing the distribution of variability of all expressed genes in the F1 blastocyst transcriptomic profiles of FVB males across the three paternal treatments. P-value computed using ANOVA. (C-D) (top) Scatter plots contrasting the variability of genes in the nABX-derived blastocysts (C) and LPHS-derived blastocysts (D) relative to controls. Genes showing higher variability in blastocysts of treated males are highlighted. (C) Genes with nABX variability – CON variability > 0.5 are shown with bigger dots. Genes with nABX variability – CON variability > 1 are shown in smaller dots. (D) Genes with LPHS variability – CON variability > 0.5 are represented with bigger dots. Genes with LPHS variability – CON variability > 1 are represented with smaller dots. (bottom) Bar chart depicting selected significantly enriched GO terms upon overrepresentation analysis of all highlighted genes in nABX (C) and LPHS (D) paternal conditions. (E) Violin and box plots comparing the distribution of variability of highly variable genes (*see* Methods) in F1 blastocyst transcriptomic profiles between the FVB and B6J backgrounds derived from control (left), nABX-treated (middle) and LPHS-treated (right) males. P-values were computed using a two-tailed unpaired t-test. (F-G) Scatter plots contrasting the variability of genes in FVB and B6 backgrounds in F1 blastocyst transcriptomic profiles from nABX-treated (F) and LPHS-treated (G) males. Genes showing high variability (Variability > 1) specific to paternal treatment in both genetic backgrounds are highlighted in brown.

To investigate whether we could capture intergenerational effects in B6 blastocysts, we elected to use ageing as a stronger environmental factor. Mammalian sperm accumulate (epi)genetic changes with age, which is linked with altered offspring phenotypes (Ashapkin *et al*, 2023; Oluwayiose *et al*, 2021). We used mature (12-13month old) B6 males – approximating middle age in humans -as sperm donors to capture F_1_ effects relative to young (9wk) males. PCA on single-blastocyst transcriptomes revealed a distinction between embryos sired by mature or young males along the second principal component (PC2) (**Fig 4E**). Moreover, we observed substantial DEGs in blastocysts derived from mature sperm donors (**Fig 4F; Table S4**). Downregulated genes were preferentially associated with splicing, and more so with oxidative phosphorylation, which is a hallmark of impaired metabolic plasticity that impacts preimplantation embryogenesis (Moussaieff *et al*, 2015) (**Fig 4F**). Upregulated genesets in offspring from older fathers were linked with immune-related processes. These data indicate that paternal age represents a significant environmental factor that triggers differential gene expression in F_1_ embryos. We speculate this may reflect a combination of accumulated genetic and epigenetic defects in aged sperm.

### Intrinsic population variability as a readout of intergenerational effects

The stark discordance between the FVB and B6 backgrounds in F_1_ responses to paternal nABX/LPHS conditioning was unexpected. We hypothesized this GxE effect may stem from intrinsically different ‘noise’ levels in blastocyst gene expression between the two genetic backgrounds. Noisy gene expression in a dynamic model obscures real signal, making it difficult to observe a consistent gene expression phenotype, which is the basis of conventional DE analysis. We therefore next asked a complementary question, namely whether paternal nABX or LPHS trigger increased variability in offspring gene activity across individual embryos. Indeed, the postnatal growth phenotype induced by paternal nABX arises probabilistically, and thus manifests as increased variation. To investigate this possibility here, we fitted a generalized linear model to the mean-variance relationship of gene expression in our data, and computed variability of a gene as the log-transformed ratio of its observed to expected variation (Fig 5A and S5A).

We first probed our FVB dataset and observed a significant difference in overall noise of blastocyst gene expression across paternal exposures (*p=0*.*024*) (**Fig 5B**). To isolate the source of this, we identified genes that exhibit substantially higher variability (ΔVariability_nABX -CON_ > 0.5 or ΔVariability_LPHS -CON_ > 0.5) in response to specific paternal perturbations, finding 292 in nABX and 438 in LPHS (**Fig 5C-D; Table S5**). Hypervariable genes showed enrichment for essential pathways like translation elongation and embryo implantation (nABX), and mRNA metabolic processes (LPHS). Amongst variable loci, we noted that MERVL retrotransposons scored as the most hypervariable across all genes and TEs specifically in response to paternal LPHS (**Fig 5D**). MERVL and its target genes have previously been implicated in the intergenerational effects of low protein diet (Boskovic *et al*, 2020; Sharma *et al*., 2016). Indeed, no MERVL effect was observed in nABX embryos, underscoring its specificity to paternal LPHS treatment. Importantly, we observed B6 blastocysts also exhibited altered gene variability in response to paternal conditioning, which included hypervariable MERVL specifically from LPHS fathers (Fig S5B-D). These data imply paternal conditioning across independent backgrounds manifests as changes in the variability of offspring gene expression profiles. High variability in essential pathways is detrimental for the success of the population, and may contribute to previous observations of disease susceptibility (Argaw-Denboba *et al*., 2024; Dalgaard *et al*, 2016).

We next compared variability between FVB and B6 background, first asking whether blastocysts exhibit equivalent baseline variation. Gene expression variability from both genetic backgrounds revealed that B6 blastocysts derived from control males are significantly noisier than their control FVB counterparts (**Fig 5E and S5E**). This implies that B6 genetics is inherently linked to more variation in expression landscapes during early development. We further found that noise in blastocysts derived from nABX or LHPS fathers is significantly greater in B6 background than FVB (**Fig 5E**). This increased gene expression noise could explain the difficulty in capturing consistent F_1_ transcriptomic responses to our paternal treatment paradigms in the B6 background.

Finally, we used gene expression variability as a score to identify genes that exhibit reproducibly elevated F_1_ variability in response to paternal perturbations across genetic contexts. We identified a signature of genes that were hypervariable in both FVB and B6 backgrounds, specifically in blastocysts from dysbiotic nABX fathers. These included genes such as *Hand1* and were significantly enriched with gene ontology terms including ‘embryonic organ morphogenesis’ and ‘response to external stimulus’, which is complementary to nABX-specific DEGs (**Fig 5F**). We further identified a set of hypervariable genes triggered specifically by LPHS-treated fathers across both B6 and FVB backgrounds. These included 2-cell stage genes *Tcstv3* and *Mervl*, and were overall enriched in ‘regulation of endocrine processes’ (**Fig 5G**). We conclude that gene expression variability is a relevant readout to capture intergenerational responses, and may represent a mechanistic route through which paternal exposures present in early embryogenesis.

### Paternal reproductive responses across environmental exposures

Given that our environmental perturbation paradigms introduce detectable F1 signatures, we finally asked how they affect the upstream molecular properties of the reproductive system in exposed fathers. We profiled the testis transcriptome, and noted that responses were driven primarily by genetic background. For example, nABX-induced dysbiosis was associated with geneset enrichment for ‘steroidogenesis’ and ‘fatty acid metabolic’ genes in FVB testes, consistent with altered lipid profiles in testis of nABX-treated males (**Fig 6A-B**). In contrast, B6 were more closely linked with immune-associated processes (**Fig 6C-D**), suggesting there is a subtle response to environmental exposures at the level of bulk transcriptome (**Fig 6C and S6A**). We also profiled testes of aged males, which elicited wholesale changes in gene expression. The majority of the DEGs in mature testes were upregulated, and corresponded to immune activation, implying increased inflammatory responses in the aged reproductive system (**Fig 6E**).

**Figure 6.**
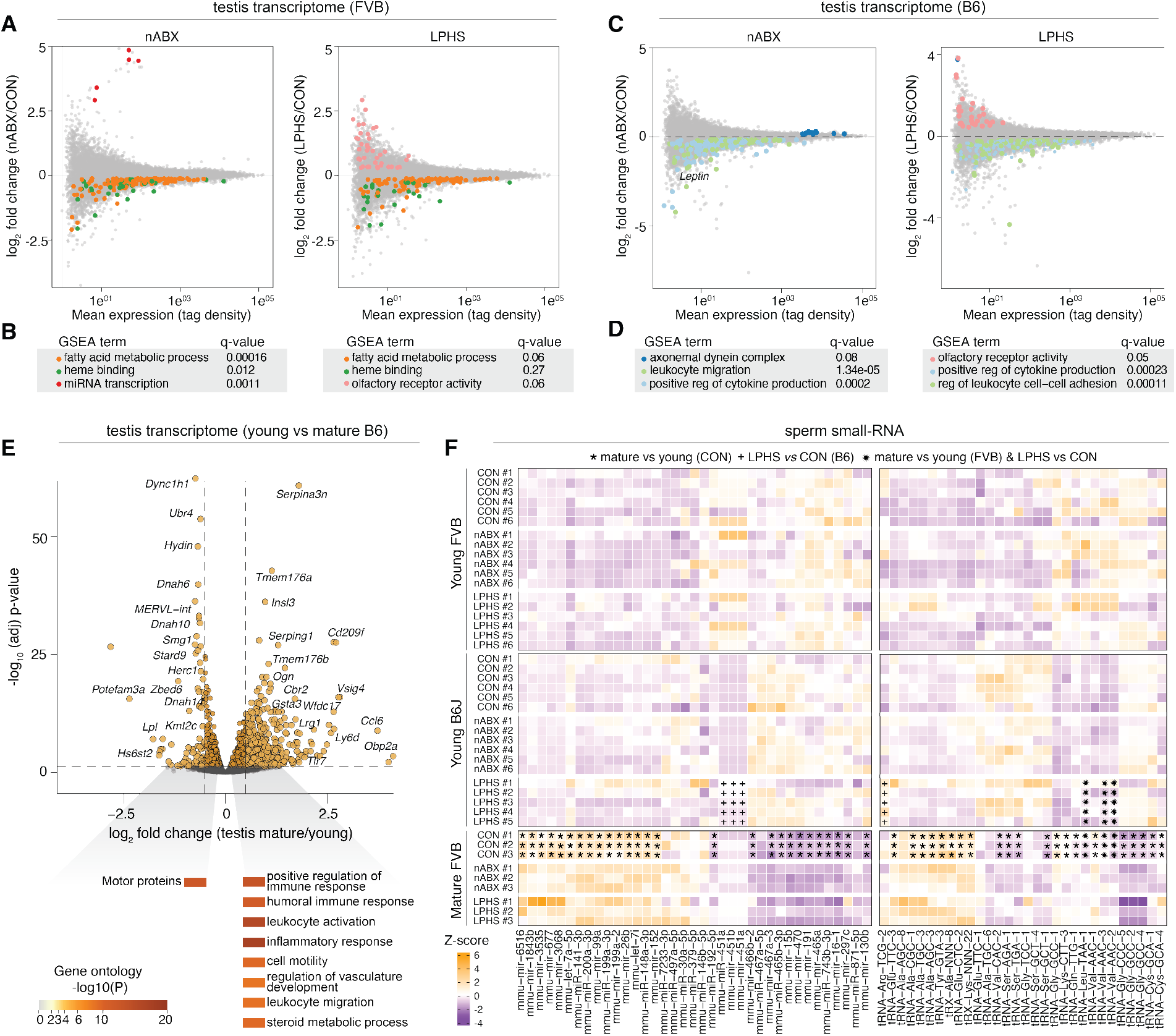
Paternal reproductive responses to environmental exposures. A-B) (A) MA plots of (left) nABX and (right) LPHS testes of FVB males versus controls highlighting genes involved in (B) enriched GSEA terms with their respective q-values. (C-D) (C) MA plots of (left) nABX and (right) LPHS testes of B6 males versus controls highlighting genes involved in (D) enriched GSEA terms with their respective q-values. (E) Volcano plot showing differential gene expression between testes of young and mature FVB males; (bottom) deconstructed heatmap highlighting select enriched GO terms in DE genes. (F) Heatmap showing relative expression of selected smallRNAs across paternal treatments, ages and genetic backgrounds. Statistical significance determined using DESeq2 adjusted p-value < 0.05. [*: Significant DE small RNA in CON mature FVB versus CON young FVB; +: Significant DE small RNA in young B6J LPHS versus CON; *: Significant DE small RNA in CON mature FVB versus CON young FVB AND in young B6J LPHS versus CON]

Multiple paradigms of intergenerational inheritance have reported differential abundance of small RNAs in sperm in response to environmental exposures (Argaw-Denboba *et al*., 2024; Chen *et al*., 2016; Gapp *et al*, 2014; Sharma *et al*., 2016; Tomar *et al*., 2024), which are proposed to partly or wholly underpin phenotypic transmission. To evaluate the signature of small RNAs in our perturbations, we adapted RNA-isolation methods in order to sequence sperm-borne small RNA from individual males (as opposed to several males pooled), and used sufficient replicate numbers (n=5-6 per condition) to ensure robustness. We confirmed that, from individual male sperm samples, we obtained the expected proportions of tRNAs, miRNAs and piRNAs, and appropriate sequencing depth (median=5.8mio post-filtering/sample) to perform differential small RNA expression analysis (**Fig S6B-C**). We observed only limited statistically significant changes in abundance of small RNA of any family in either the FVB or B6 backgrounds, in both the nABX and LPHS treatment paradigms (**Fig 6F and S6D-E**). A small number of tRNA-derived small RNAs and miRNAs were significantly abundant upon LPHS treatment in the B6 background (**Fig 6F**). Paternal age was, by some distance, the strongest environmental perturbation, with miRNAs and tRNAs showing significant expression changes in sperm from older males (**Fig 6F and S6F; Table S6**).

To further understand the limited changes in miRNA and tRNA upon paternal environmental exposure in young males, we examined expression of small RNA within each individual. We identified considerable heterogeneity in expression levels in sperm from young males (**Fig 6F**). In contrast, mature sperm samples exhibited a much more consistent expression profile across replicates (**Fig 6F**). These results imply an inherent variability in small RNA levels in sperm of younger males, which are typically deployed in intergenerational studies. Taken together, these data indicate that sperm-borne small RNA do not exhibit consistent and significant environmental responses when using relatively high-powered experimental designs, while paternal age acts as a strong modulator of the reproductive system transcriptome and sperm small RNA payloads. This consideration should be taken into account when dissecting the mechanisms of transmission for intergenerational phenotypes and indeed, the triggers of the F1 molecular signatures reported in the present study.

## Discussion

In this study, we executed a highly controlled strategy with large replicate numbers to comprehensively characterise the earliest molecular signatures of mammalian paternal effects. Across all IVF experiments, we generated transcriptomic profiles from 808 individual embryos spanning three paternal treatments, two paternal ages and two genetic backgrounds. The results reveal a reproducible transcriptomic signature of the preconception paternal condition emerges in F_1_ blastocysts. The data further show that such signatures are typically subtle, and interact with underlying genetics, age, and potentially additional factors to greatly influence overall outcome. This study therefore reinforces that transmission of paternal environmental factors constitutes an important source of F_1_ phenotypic variation, but also that complex and confounding influences play into the penetrance and detection of such effects.

One intriguing observation is that nABX-induced paternal gut microbiome dysbiosis may mechanistically operate by influencing early lineages in embryos. We found that across independent experimental batches, paternal dysbiosis led to reduced expression of extra-embryonic lineage markers, particularly primitive endoderm (exemplified by *Gata4)*, with concurrent higher expression of epiblast lineage genes. This could reflect an altered ratio of (extra-)embryonic lineage allocation and/or specific changes in the expression of master regulators within lineages. In principle, even a modest imbalance in lineage identity and/or embryo size could propagate through development and elicit a temporally-uncoupled emergent phenotype in mature tissues, such as placenta (Greenberg *et al*, 2017; Perez-Garcia *et al*, 2018; Simon *et al*, 2018). Indeed, paternal dysbiosis has been shown to affect F1 phenotype probabilistically, via impacting placental development (Argaw-Denboba *et al*., 2024). In contrast to nABX, we found paternal exposure to suboptimal low-protein, high-sugar (LPHS) diet could be linked with a modest developmental delay, as judged by smaller embryos and reduced expression of ribosomal/MAPK genes, which regulate proliferation. Such effects of paternal diet on embryonic growth/proliferation have been noted before, as have MERVL responses (Sharma *et al*., 2016), and can lead to impaired implantation (Modzelewski *et al*., 2021). Taken together these observations imply that that distinct paternal environments elicit relatively specific effects on progeny, at least for the initial presentation.

Our study also marks a note of caution in interpreting the impact of intergenerational effects. For example, we found genetic background fundamentally affects the prevalence and signature of paternal effects. This implies that disentangling GxE is not only key to understand phenotypic traits in individuals, but also for their propagation to the next generation (intergenerational GxE or iGxE). The extent to which the difference between backgrounds represents biology or technical challenges is unclear however. Indeed, B6 embryos are inherently more variable in their gene expression and developmental success rates, which impairs the ability to detect consistent and subtle paternal effects. Conversely, we propose that the use of noise (or excess variability) actually represents an important phenotyping strategy. Paternally-induced high embryo-to-embryo variation in expression was prevalent across our paradigms, and could help explain observations of partial penetrance in intergenerational studies. We also envision that deploying other modalities as readouts of early F_1_ phenotypes, such metabolomics or proteomics, will capture a fuller spectrum of F_1_ effects. Finally, our data indicates it is critically important to ensure intergenerational readouts are sufficiently powered, especially given the often subtle impact of paternal effects. Smaller-scale experiments run a risk of under-sampling the distribution of molecular/phenotypic outcomes, and consequently over-or under-estimating penetrance and effect-size.

Thus, we propose some recommendations that could strengthen the conclusions of intergenerational studies: (i) it is important to profile high numbers of F_1_ embryos/progeny, across multiple experimental batches with multiple fathers, to maximise signal-to-noise ratio, (ii) examining several genetic backgrounds reveals both biological insights for iGxE and generalisability of transmitted effects, and (iii) partial phenotypic penetrance (or excess variability) could serve as an important strategy for observing intergenerational effects. In conclusion, our study provides evidence for a molecular signature of paternal environmental exposures at the earliest stages of offspring development, and provides a template for future studies to achieve a mechanistic understanding of mammalian intergenerational epigenetic inheritance.

## Acknowledgements

Funding for this study was supported by the EMBL Human Ecosystems Transversal Theme (59100) and programme funding to J.A.H and A.B. We thank the GEVF and GBCS core facilities for experimental and technical support.

## Author contributions

CRediT attribution. M.D: Investigation (lead), Formal analysis, Visualisation, Writing – original draft, review & editing; B.R: Formal analysis (lead), Data Curation, Writing -original draft, review & editing, Visualisation; V.P: Investigation; R.P: Investigation; J.S: Investigation: L.V: Investigation; V.B: Resources; O.B: Resources; A.B: Conceptualisation (equal), Funding acquisition, Writing – review & editing, Supervision; J.A.H: Conceptualisation (equal), Formal analysis, Visualisation, Funding acquisition, Writing – original draft, review & editing (lead), Supervision (lead).

## Data availability

All data supporting the findings of this study are available and have been deposited into ArrayExpress under accessions: E-MTAB-14640, E-MTAB-14638, E-MTAB-14639, E-MTAB-14635, E-MTAB-14636, E-MTAB-14637.

## Code availability

All code used for data analysis in this study is available on GitLab at: https://git.embl.de/grp-hackett/iei-transcriptomics

## Materials and Methods

### Section I: Exposures and embryology

#### Animal husbandry

All experiments involving mice were carried out in accordance with the approved protocols and guidelines by the laboratory animal management and ethics committee of the European Molecular Biology Laboratory (EMBL) under license no. 22017RMJH. The inbred FVB, C57BL/6J (B6), and C57BL/6NJ (B6) strains were used in this study. Mice were housed under a 12 h light/dark cycle (light from 07:00 to 19:00), with *ad libitum* access to food and beverage (see detail in the *“mice environmental treatment”* methods section). Mature male mice (1 year) were kept as much as possible with their littermates in the same cage but isolated upon signs of aggression. Animals were sacrificed by cervical dislocation for all experiments.

#### Mice environmental treatment

At the onset of each treatment, male mice were randomly allocated to one of three groups: Control (CON), non-absorbable antibiotics (nABX), and low protein high sugar diet (LPHS). Each group received the appropriate exposure regimen after 3 weeks or 12-13 months, respectively, for young or mature mice (Fig. 1A and 4A). CON cages were supplied with water and a rodent standard diet (protein 18,5%, sucrose 3,7%, see extensive Mucedola composition in Table S7), nABX cages received non-absorbable antibiotics in drinking water (Neomycin trisulfate salt hydrate (Sigma N1876) (final concentration: 2.5mg/mL), Bacitracin (Sigma B0125) (final concentration: 2.5mg/mL), and Pimaricin (Sigma P9703) (final concentration: 1.25μg/mL)), in addition to the standard diet (composition provided above). Finally, LPHS cages were provided with water and a western-style, low-protein high-sugar (LPHS) diet (cornstarch 39.75%, sucrose 19.1%, casein 10.9%, maltodextrin 13.2%, soybean oil 7%, cellulose 5%, mineral mix 3.5%, vitamin mix 1%, L-Cystine 0.3%, choline bitartrate 0.25%, tert-butyl hydroquinone 0.00%, ref: Carone, Cell, 2010). Mice were monitored daily for well-being and weighed weekly. Following 6-7 weeks of continuous treatment, mice were collected for downstream experiments.

#### In vitro fertilization (IVF)

*In vitro* fertilizations were performed based on published protocols, with some modifications to optimise throughput and equalise developmental timing (Guan *et al*, 2014). Briefly, cauda epididymis and vasa deferentia from males were dissected and sperm was gently squeezed/dissected out into preequilibrated capacitation media (HTF supplemented with MBCD at final concentration of 0.75 mM) under mineral oil (HTF: Millipore MR-070-D; MBCD: Sigma-Aldrich C4555; mineral oil: sigma M8410, need to be batch tested), and allowed to swim up for 1 hour. In the meantime, cumulus-oocyte complexes were collected from superovulated females in the fertilization drop containing KSOM media with GSH (final concentration 1mM) under mineral oil (KSOM: Millipore MR-106-D; GSH: L-Glutathione reduced, Sigma G6013). Superovulated females (6-12 weeks for FVB and 3-4 weeks for B6) were intraperitoneal injected with 5-10 IU of PMSG and 5-10 IU of hCG 64h and 16h before oocyte collection, respectively. Then, sperm were counted, and cumulus-oocyte complexes were inseminated with 0.2 million sperm in a 200 μl fertilization drop (KSOM + GSH). Co-incubator proceeded for four hours in a low oxygen condition (hypoxic) incubator (5% CO_2_, 5% O_2_, 37°C). Following co-incubation, zygotes were cleaned from the surrounding cumulus cells and sperm by 5-8 washes in KSOM covered by mineral oil, then cultured in KSOM in a 96-well plate (without mineral oil) until the blastocyst stage (92h +/-2h after completed fertilization) in low oxygen condition incubator (5% CO_2_, 5% O_2_, 37°C) or regular condition incubator (5% CO_2_, 20% O_2_, 37°C). As a side note, the genetic background of the female always matched the genetic background of the male (e.g. for the paternal condition involving mature FVB, *IVF* was conducted using FVB oocytes).

#### Blastocyst analysis with COPAS vision

FVB control and treated blastocysts were generated by *IVF* (as described) and cultured in a normoxic incubator (5% CO_2_, 20% O_2_, 37°C). Blastocysts were washed in 1X PBS, fixed in 4% PFA for 20min at 37°C (with their zona pellucida), and transferred in a 50mL falcon tube containing 20mL of 1X PBS. Then, blastocysts were resuspended in solution by vigorously shaking the falcon tube immediately before starting the analysis by the COPAS vision. Analyses were conducted with a flow cell of 500μm, and 8-bit images were captured for each object. Blastocysts were characterized based on Time Of Flight (TOF) and Extinction (optical density, meaning light reflection capacity). The COPAS vision software (FlowPilot 3.091) automatically computed various morphological features of the blastocyst embryo, including perimeter in μm, area in μm2, roughness in arbitrary unit (A.U.) (roughness represents the deviation from 1 with 1 being completely round); and mean grayscale in A.U. (for 8-bit images, grayscale should be an integer value between 0 and 255 representing the gray level of the pixel with 0 being black and 255 white).

### Section II: Omics

#### Single embryo RNA-sequencing

Single-embryo RNA-sequencing was performed based on the previously published SMART-Seq protocol (Hennig *et al*, 2018; Picelli *et al*, 2014; Trombetta *et al*, 2014) with some modifications outlined below.

#### Embryo collection for SmartSeq

Blastocysts were generated from *IVF* and cultured in a hypoxic environment (5% CO_2_, 5% O_2_, 37°C) for all experiments and conditions, except for comparison of young and mature C57BL/6N (B6) paternal states (Fig 4E-F), where normoxic incubation was performed (5% CO_2_, 20% O_2_, 37°C). Embryos were washed 2-3 times in warm M2 media (Sigma M7167), and individually mouth pipetted into 5uL of collection buffer (Trition-X-100 0,5%; RNase OUT 0,5U/uL - Invitrogen 10777-019; dNTPs 2.5mM and oligo-dT 2.5uM (5’AAGCAGTGGTATCAACGCAGAGTACTTTTTTTTTTTTTTTTTTTTTTTTTTTTTTVN 3’) in RNAse free water. Embryos were stored in the collection buffer at -80°C prior to library preparation.

#### SMART-Seq library preparation

*(protocol 1; all data except Fig 4E-F)*. SMARTseq library preparation was performed following these steps. The samples were organized into a skirted 96-well plate (Fisher Scientific E951020401), incubated for 3 minutes at 72°C, immediately placed on ice, and then centrifuged. The first step was the reverse transcription of mRNA. An RT master mix (final volume: 6.15 μL) was prepared with the following components: 2 μL 5X SuperScript IV buffer (Thermo Scientific 18091050), 0.50 μL DTT 100mM, 2 μL Betaine 5M (Sigma B03000 1VL), 0.1 μL MgCl2 1M, 0.25 uL RNAse inhibitor (Clontech Takara 2313A), 0.25 μL SuperScript IV RT 200U/μL (Thermo Scientific 18091050), 0.1 μL template switching oligo 100μM (5′-AAGCAGTGGTATCAACGCAGAGTACATrGrG+G-3′), and 1.15 uL of water. This master mix was added to each well, containing a single embryo in 5 μL of collection buffer. Each well was thoroughly mixed by pipetting, sealed, centrifuged, and incubated in a thermocycler under the following conditions: 42°C for 15min, 80°C for 10min, then stored at 10°C.

The second step, involving whole transcript amplification (WTA) and post-PCR cleanup, was immediately performed. The plate was unsealed, and a WTA mix was added to each sample (final volume: 15 μL; 12.5 μL KAPA HiFi HotStart ReadyMix 2x (Roche); 0.20 μL IS PCR Primer 5μM (5′-AAGCAGTGGTATCAACGCAGAGT-3′), and 2.30 uL water). The samples were mixed well by pipetting, sealed, and incubated in a thermocycler under the following conditions - Initial step: 98°C for 3min; 17-18 cycles: 98°C for 20sec, 67°C for 15sec, 72°C for 6min; Extension: 72°C for 5min; and then storage at 10°C. The PCR products were then cleaned up using DNA SPRI beads (Beckman Coulter) at a 0.6 volume ratio. The mixture was incubated 5min at RT with the beads, separated using a magnet, and immediately eluted in 13uL of water. As a note, the usual ethanol bead wash steps were not necessary since the cDNA was highly diluted, and washing seemed to decrease the cDNA yield. The cDNA concentration and profile were verified using an HS DNA Bioanalyzer chip. For the young FVB paternal condition, sample from the same 96-well plate were pooled and run only once in the Bioanalyzer chip. The median concentration was calculated and applied for later dilution of all samples from this plate. For the mature FVB and young B6J paternal condition, each cDNA sample was quantified one by one and diluted.

During the third step, called tagmentation, homemade reagents were used in the process.

First, the linker oligonucleotides were annealed. In details, the oligos Tn5ME-A (5′-TCGTCGGCAGCGTC<u>AGATGTGTATAAGAGACAG</u>-3′), Tn5ME-B (5′-GTCTCGTGGGCTCGG<u>AGATGTGTATAAGAGACAG</u>-3′), Tn5MErev (5′-[phos]<u>CTGTCTCTTATACACATCT</u>-3′) were resuspended in an annealing buffer (50mM NaCl, 40mM Tris-HCl pH8.0); equally mixed (Tn5ME-A + Tn5MErev; or Tn5ME-B + Tn5MErev) at a final concentration of 50uM; annealed in the thermocycler for 5min at 95°C, slowly cool down to 65°C (0.1°C/sec), 5min at 65°C and slowly cool down to 4°C (0.1°C/sec); and diluted to 35uM in water.

Second, the Tn5 was loaded with the annealed linkers. Homemade Tn5_(R27S),E54K,L372P_ was diluted at 0,5mg/mL. Annealed linkers (0.7uL at 35uM) were mixed with 10uL of diluted Tn5 (0.5mg/mL), incubated at 23°C under constant shaking at 350rpm for 30-60min and store on ice. Finally, loaded Tn5 was diluted in water of a final dilution 1:200 to 1:300. Third, cDNA was diluted to 100pg/uL to 200pg/uL and 1.25uL of diluted cDNA was transferred into a new 96-well plate. Tagmentation reaction was added to each sample 2.5uL of tagmentation mix (1 volume of 4x tagmentation buffer (40 mM Tris-HCl pH 7.5, 40 mM MgCl2) and 1 volume of 100% DMF (Sigma Aldrich)), 1.25uL of Tn5 (1:200 to 1:300), mixed well (final volume 5uL); incubated at 55°C for 3min and place immediately on ice. The reactions were stopped by adding 1.25uL of 0.2% SDS, incubated 5min at RT and stored on ice.

Final PCR was performed. In details, 10uL of the following PCR mix was added to each samples (6.75uL of 2x KAPA buffer (Roche), 0.75uL of 100% DMSO, 1.25uL of 10uM i5 adapter index primer, and 1.25uL of 10uM i7 adapter index primer) and incubated in the thermocycler under the following conditions: 72°C for 3min; 95°C for 30sec; 12 cycles: 98°C for 20°C, 58°C for 15sec, 72°C for 30sec; 72°C for 3min; and store at 10°C. Then, all samples were pooled (5-8uL / sample), and the final pool was cleaned with 0.8x SPRI beads and eluted in 12uL. The concentration and quality of the pooled library were assessed using the Agilent HS DNA BioAnalyzer D1000 and the Qubit dsDNA HS assay.

#### SMART-Seq library preparation

*(protocol 2) – (Fig 4E-F)*. Young and mature B6N paternal condition SMARTseq library preparation was made only using commercial reagents. Briefly, the samples were organized into a skirted 96-well plate (Fisher Scientific E951020401), incubated for 3 minutes at 72°C, immediately placed on ice, and then centrifuged (all centrifugation steps were done at 800g for 1 minute). The first step was the reverse transcription of mRNA. An RT master mix (final volume: 6.65 μL) was prepared with the following components: 2 μL Betaine 5M (Sigma B03000 1VL), 2 μL 5X SuperScript IV buffer (Thermo Scientific 18090050), 0.25 μL DTT 100mM, 0.9 μL MgCl2, 1 μL template switching oligo 10μM (5′-AAGCAGTGGTATCAACGCAGAGTACATrGrG+G-3′), and 0.5 μL SuperScript IV RT 200U/μL (Thermo Scientific 18090050). This master mix was added to each well, which already contained a single embryo in 5 μL of collection buffer. Each well was thoroughly mixed by pipetting up and down, sealed, centrifuged, and incubated in a thermocycler under the following conditions - initial step: 42°C for 90 minutes; 10 cycles: 50°C for 2 minutes and 42°C for 2 minutes; inactivation: 70°C for 15 minutes; and then stored at 4°C.

Next, the second step, which involved whole transcript amplification (WTA) and post-PCR cleanup, was performed. The plate was unsealed, and a WTA mix was added to each sample (final volume: 14 μL; 13 μL KAPA HiFi HotStart ReadyMix (Thermo Scientific 18090050); 1 μL IS PCR Primer 10μM (5′-AAGCAGTGGTATCAACGCAGAGT-3′)). The samples were mixed well by pipetting, sealed, and incubated in a thermocycler under the following conditions - Initial step: 98°C 3min; 20 cycles: 98°C 15sec, 67°C 20sec, 72°C 6min; Extension: 72°C 5min; and then storage at 4°C. The PCR products were then cleaned up using DNA SPRI beads (Beckman Coulter B23318) at a 0.8 volume ratio with 80% EtOH. The elution step was performed in 20 μL of clean water, and the eluents were transferred to a new skirted 96-well plate. Each sample was quantified using the Qubit dsDNA HS Assay Kit with Qubit assay tubes and then diluted to 0.5 ng/μL in clean water in a new skirted 96-well plate.

The third step involved constructing the Nextera XT sequencing library (Illumina 15032354). Firstly, the i5 and i7 primers were combined in a 96-well plate as detailed in the commercial protocol. Secondly, the tagmentation reaction was performed. In a new skirted 96-well plate, the tagmentation mix (2.5 μL TD buffer and 1.25 μL ATM) was combined with 1.25 μL of diluted product from samples (concentration 0.5 ng/μL) and mixed well. Tagmentation was carried out in a thermocycler for 10 minutes at 55°C and then cooled down to 10°C. As soon as the tagmentation reaction reached 10°C, 1.25 μL of NT buffer was added to each well and incubated for 5 minutes at room temperature to stop the reaction. Thirdly, indexes were incorporated (Nextera XT Index Kit v2 Set B, REF 15052164, LOT 20756419), and the products were amplified. In detail, each sample received 3.75 μL of NPM and 2.5 μL of the specific combined index primer solution. The samples were mixed well by pipetting, centrifuged, and incubated in a thermocycler with the following program - initial annealing: 72°C 3min; denaturation: 95°C 30sec; 12 cycles: 95°C 10sec, 50°C 30sec, 72°C 1min; extension: 72°C 5min; and then storage at 4°C.The final step is the DNA pooling and cleanup. Briefly, 2.5 μL of each sample was pooled into a 1.5 mL Eppendorf tube, washed with DNA SPRI select beads at a 0.9 volume ratio with 80% EtOH, and then eluted in 30 μL of clean water. This washing step was repeated again with DNA SPRI select beads at a 0.9 volume ratio with 80% EtOH, and the DNA was eluted in 30 μL of clean water. Finally, the quality and the concentration of the pooled library was assessed using the Agilent HS DNA BioAnalyzer D1000 and the Qubit dsDNA HS assay.

#### SmartSeq library sequencing

After SMART-seq library preparation of single embryos, the indexed pooled samples were sequenced together in one or two runs using the NextSeq 2000 sequencer with a P3 flow cell in 50bp paired-end mode. Numbers of obtained reads after QC are available in Supp. Fig. 1J, 4aG.

#### Testis RNA-seq

Whole testes from FVB, C57BL/6J control and treated mice were collected, snap-frozen in liquid nitrogen, and stored at -80°C. Testes were then homogenized in Trizol using a BeadBug^™^ 6 microtube homogenizer with 1mm glass beads (Accessories Beads for BeadBeater® Zirconia/glass beads, 0,1 mm – Roth No. N033.1). Total RNA was extracted according to the manufacturer’s recommendations. RNA was quantified using Nanodrop and RNA TapeStation kit. RNA-seq libraries were performed with a starting material ranging from of 100-500ng/uL of total RNA using the NEBNext Ultra II directional RNA kit with the Beckman Coulter hybrid workstation i7 (liquid handling robot). Adaptors were diluted 1 in 30. Libraries were amplified through 11 PCR cycles and sequenced with a 40bp, pair-ends reading mode on a NextSeq500 sequencer.

#### Sperm small RNA-seq

##### Sperm collection

The cauda epididymides from FVB, C57BL/6J control and treated male mice were dissected and placed into a 1.5mL Eppendorf tube containing warm M2 media (Sigma M7167). The caudae and M2 media were then transferred into a 6cm dish and gently cleaned from any surrounding tissue or fat. After cleaning, the caudae were briefly washed in clean M2 and placed in a 3.5cm dish containing 1-1.5mL of M2 media. The caudae were then cut open approximately 20 times with scissors and incubated for 30min at 37°C, 5% CO2, to allow spermatozoa to swim out. Under the binocular loupe, the areas of tissue still containing spermatozoa were identified and pinched with needles to release the remaining spermatozoa and incubated for an additional 10min at 37°C, 5% CO2. Following incubation, the M2 media containing the spermatozoa was pipetted up and down a few times and transferred into a fresh 1.5mL Eppendorf. It is important to perform this procedure carefully to avoid excessive disruption of the tissue and, therefore, prevent somatic cell contamination of the sperm collection. The sperm solution was filtered through a 40uM filter to remove any tissue fragments. The filtered solution was then centrifuged at 8500rpm, 5min, 4°C. The pellet was resuspended in 1mL of somatic cell lysis buffer (0.1% SDS, 0.5% Triton-X-100, in water) and incubated on ice for 20min. Following somatic cell lysis, the sperm solution was pelleted (first at 8500rpm and then at 13500rpm, both for 5min at 4°C) and resuspended in 1mL of 1X PBS each time. Finally, the spermatozoa were pelleted again (13500rpm, 5min, 4°C), quickly dried, and frozen at -80°C for storage.

##### Sperm total RNA extraction

Sperm RNA extraction was performed using an optimized version of mirVana miRNA isolation kit (Thermo fisher scientific AM1560). Spermatozoa were first resuspended in 600 uL of mirVana lysis buffer and transferred into a screw-cap tube containing 1mm beads (Accessories Beads for BeadBeater® Zirconia/glass beads, 0,1 mm – Roth No. N033.1). The spermatozoa were homogenized for 60sec at 4000rpm for a total of 5 cycles using the BeadBug™ 6 microtube homogenizer. Samples were centrifuged at RT, 5000rpm, 5min and cooled on ice for 10min to complete sperm lysis. Avoiding bead collection, the solution was transferred to a new 1.5mL Eppendorf tube. Following this, the “organic extraction” and “Total RNA isolation procedure” sections from the mirVana protocol were precisely followed. Briefly, the extracted RNA was incubated on ice for 10min with 1/10 volume of miRNA Homogenate Additive and isolated with an Acid-Phenol: Chloroform step. The aqueous phase containing the RNA was recovered and mixed with 1.25 volume of 100% RT ethanol. The lysate/ethanol mixtures were applied to a filter cartridge and washed 3 times. RNA was eluted with 50uL of pre-heated nuclease-free water (95°C). RNA concentration and profile were measured using the RNA High Sensitivity TapeStation kit (measured RNA concentration between 1000 and 8000 pg/uL). Sperm RNA can be stored at -80°C for weeks to months.

##### Small RNA library construction and sequencing

Libraries were constructed using the NextFlex Small RNA-Seq Kit v4 by PerkinElmer. The starting material for the libraries varied in concentration for each sample, while the volume remained consistent across all samples. Libraries were sequenced at 100bp, single end using a Nextseq2000 sequencer, yielding between 100,000 and 7 million reads per sample, with the vast majority of the sample containing approximately 1-5 million reads.

### Section III: Single-embryo transcriptomics analysis

#### Constructing the reference genome

We used STAR (version 2.7.11a) *genomeGenerate* function to modify the default mm39 genome (which is based on a B6 background) by adding an FVB-specific SNP mask obtained from dbSNP^1^ lifted over to the mm39 genome. To map transcriptomic sequences obtained from B6J and B6N backgrounds, the mm39 genome was used as is. TE annotations were obtained from UCSC RepeatMasker and added to the respective genome’s GTF file prior to running *genomeGenerate*. Key redefined parameters:

- --genomeTransformType Haploid
- --sjdbOverhang 99
- --genomeChrBinNbits 15

#### Mapping and alignment of single-embryo transcriptomics data

Raw reads were mapped to the FVB SNP-masked or standard mm39 genome with TE annotations using STAR (version 2.7.11a). To include transposable elements in our dataset, we followed recommendations from Teissandier et al., 20192. Briefly, we allowed for multi-mapping up to a maximum of 5000 candidate loci per read and randomly assigned the mapped read to any of its candidate loci. Key redefined parameters:

- --outFilterMultimapNmax 5000
- --outFilterMismatchNoverLmax 0.06
- --winAnchorMultimapNmax 5000
- --outSAMmultNmax 1

#### Quantification of reads from genes and transposable elements

Read quantification was performed using featureCounts (v2.0.6) separately for genes (at default settings) and transposable elements (with “-M” parameter to include multi-mapped reads).

#### Principal Component Analysis (PCA)

All downstream analysis was performed using R (4.2.2). Briefly, counts were normalised to counts-per-million (CPM) using edgeR (3.40.2) package’s *cpm* function and subsequently log_2_-transformed with a pseudocount of 1. Log-normalised counts were then input into the *prcomp* function of the irlba (2.3.5.1) package at default parameters. Density plots lining the PCA plots were generated using the *geom_density* function of ggplot2 (3.5.1).

#### Differential expression analysis

Differential expression analysis was performed using DESeq2 (1.38.3) (see GitLab link for full code and parameters). Genes showing an adjusted p-value < 0.05 and log2FoldChange > 0.5 were identified as DE genes in all cases. An exception was applied specifically when comparing young vs mature F1 blastocysts (B6), where we highlighted genes with adjusted p-value < 0.05 but no wit log2FoldChange threshold ino order to capture highly significant but low effect-size hits. Overrepresentation analysis to obtain Gene Ontology terms amongst DEG (Fig 1G-H) was performed using String v12.0.

#### Clustering of DE gene sets

Expression of DE genes obtained from both nABX and LPHS conditions in the young FVB background was averaged over all samples in the same condition. For each gene, the averaged expression in each condition was then scaled by its average expression in Ctrl samples. We then constructed a gene-gene euclidean distance matrix using *dist* with *method = “euclidean”*, and clustered the distance matrix by complete linkage hierarchical clustering using *hclust* with *method = “complete”*. The *cutree* method was used to cut the dendrogram and obtain distinct clusters, which were then manually merged based on directionality of expression relative to controls to obtain the 8 clusters in Figure 1I. Overrepresentation analysis to identify enriched Gene Ontology (GO) terms in DE gene sets was performed using Metascape (v3.5.20240101) “Express Analysis” mode at default settings3.

#### Gene Set Enrichment Analysis (GSEA)

GSEA was performed using clusterProfiler (4.6.2). Briefly, genes were ordered based on log2FoldChange output from the differential expression analysis above with *nPermSimple=10000 & pAdjustMethod = “BH”*.

#### Construction of single-cell reference and computing correlations

Single-cell transcriptomic profiles were downloaded from Deng et al., 2014^4^. Specifically, mid blastocyst (GSM1112706 - GSM1112765) and late blastocyst (GSM1112664 - GSM1112693) profiles were downloaded and subset for expression of lineage markers. Lineage marker expression profiles of single-cells were centered and scaled, then used to compute a cell-cell distance matrix, which was then clustered using *hclust*. Cells belonging to epiblast and trophectoderm lineages were obtained by cutting the dendrogram using *cutree* while primitive endoderm cells were annotated based on expression of primitive endoderm-specific markers. Single-cell expression profiles were then averaged over cells of the same cluster to obtain an average expression signature of lineage-specific marker genes for each lineage.

Expression of our single-embryo data over the lineage markers was then correlated (spearman correlation) with the average expression signatures of each lineage to obtain the correlation coefficients observed in Figure 2a.

#### Profiling binding of transcription factors in our DE gene sets

Binding of transcription factors reported across various cell and tissue types was performed using Enrichr5.

#### Remapping published ChIP-seq datasets to obtain chromatin mark profiles

ICM ChIP-seq FASTQ files were downloaded from Liu et al., 2016 (H3K4me3 & H3K27me3), Wang et al., 2018 (H3K9me3) and Xu et al., 2019 (H3K36me3). Raw reads were trimmed using TrimGalore (v0.6.10) with a quality threshold of 20 to clip the 1 base from the 5’ end of both paired reads. Trimmed reads were then mapped to the mm39 reference genome using bowtie2 (v2.4.2). Blacklist regions for mm39 were obtained from excluderanges6 and excluded prior to quantification. Duplicate reads were marked and removed using *MarkDuplicates* (v3.1.0). Finally, coverage files were generated using *bamCoverage* module of deeptools (v3.5.5) with RPKM normalisation and replicates were merged.

Enrichment of reads around the TSS of DE genes was computed using *computeMatrix reference-point* mode of deeptools with the following parameters: -*a 5000, -b 5000*, -*binSize 500*. Heatmaps and enrichment profiles seen in Figure 2d were plotted using the *plotHeatmap* module of deeptools at default settings.

### Section II: Variability analysis

#### Calculating variability of expression of a gene

Our variability analysis is inspired by work from Conine et al., 20207. For each gene, mean expression and coefficient of variation were calculated and a generalised additive model (GAM) was fit to the CV-mean distribution using the *gam* function of the R package mgcv (1.8-42) using the following parameters: *formula = CV ∼ s(Mean, bs = “cs”, fx=TRUE, k=20), fit = TRUE, method = “GCV*.*Cp”*. Expected CV was obtained using the *predict* function of R stats using the following parameters: *se = TRUE, type = “link”*. Variability of a gene was calculated using the following formula:

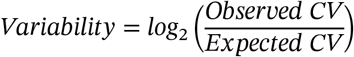

All overrepresentation analyses were performed as described above.

### Section IV: Testis transcriptomics analysis

#### Mapping and alignment of testis transcriptomics data

Testis transcriptomics data was mapped and aligned as described above for the single-embryo transcriptomic data.

#### Quantification of reads from genes and transposable elements

Read quantification was performed as for single-embryo transcriptomics data with one key difference – counting was performed in a strand-specific manner using the parameter *-s 2*.

PCA, differential expression analysis and GSEA were performed as described above for the single-embryo transcriptomic data.

### Section V: Analysis of small RNA profiles from sperm

#### Trimming, filtering, QC, mapping and alignment of small RNA sequencing data

Raw reads were trimmed using TrimGalore (v0.6.10) in smallRNA mode with quality threshold of 20 and restricting length of reads from 18 to 40 bases. Key redefined parameters:

- --small_RNA
- --quality 20
- --length 18
- --max_length 40

Trimmed reads were mapped using bowtie (v1.0.0) allowing for no mismatches and reporting only a single best alignment per read. Key redefined parameters:

- -v 0
- -k 1
- --best

#### Quantification of reads from miRNAs and tRNAs

miRNA annotations were obtained from miRBase8 and lifted over to mm39 using CrossMap (v0.7.0) at default settings. tRNA annotations for the mm39 genome were obtained from GtRNAdb9. miRNA and tRNA quantification were performed by counting intersections between the respective reference locations and locations of trimmed reads on the mm39 reference genome using bedtools (v2.31.1) *intersect* at default settings.

#### Differential expression analysis

Differential expression analysis was performed as described above for single-embryo transcriptomics data.

