## Supplemental Figs 1-6 for "Embryonic Signatures of Intergenerational Inheritance across Paternal Environments and Genetic Backgrounds"

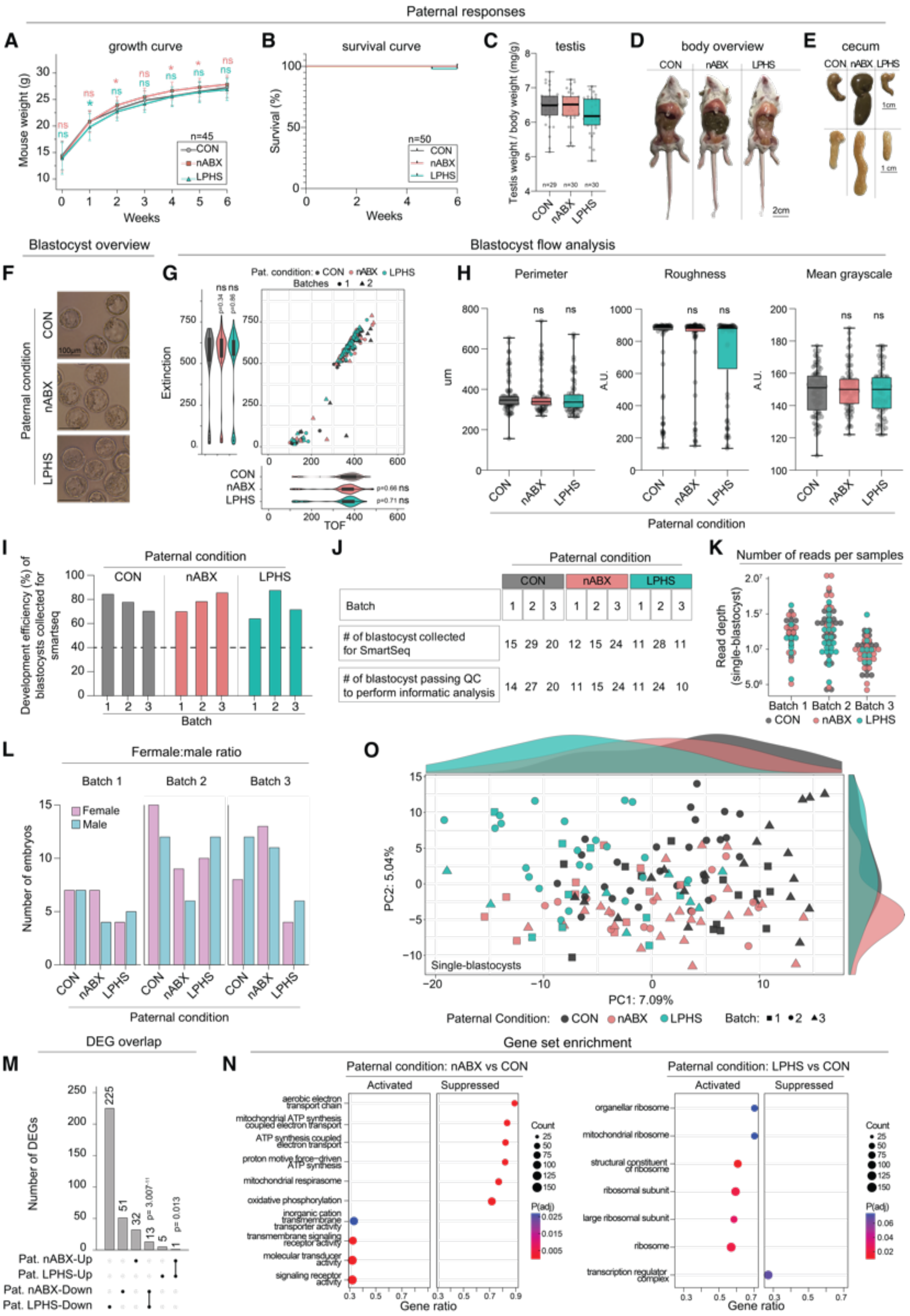
Paternal responses
A
growth curve
Mouse weight (g)
Weeks
n=45
CON
nABX
LPHS
B
survival curve
Survival (%)
Weeks
n=50
CON
nABX
LPHS
C
testis
Testis weight / body weight (mg/g)
CON
nABX
LPHS
D
body overview
CON
nABX
LPHS
2cm
E
cecum
CON nABX LPHS
1cm
Blastocyst overview
Blastocyst flow analysis
F
Paternal condition
CON
nABX
LPHS
100μm
G
Pat. condition: • CON • nABX • LPHS
Batches • 1 ▲2
Extinction
TOF
H
Perimeter
Roughness
Mean grayscale
um
A.U.
A.U.
Paternal condition
I
Paternal condition
CON
nABX
LPHS
Development efficiency (%) of blastocysts collected for smartseq
Batch
J
Paternal condition
CON
nABX
LPHS
Batch
1 2 3 1 2 3 1 2 3
### of blastocyst collected for SmartSeq
15 29 20 12 15 24 11 28 11
### of blastocyst passing QC to perform informatic analysis
14 27 20 11 15 24 11 24 10
K
Number of reads per samples
Read depth (single-blastocyst)
Batch 1 Batch 2 Batch 3
• CON • nABX • LPHS
L
Femlale:male ratio
Batch 1
Batch 2
Batch 3
Female
Male
Number of embryos
Paternal condition
O
PC2: 5.04%
PC1: 7.09%
Single-blastocysts
Paternal Condition: • CON • nABX • LPHS
Batch: ■1 •2 ▲3
DEG overlap
M
Number of DEGs
Pat. nABX-Up
Pat. LPHS-Up
Pat. nABX-Down
Pat. LPHS-Down
Gene set enrichment
N
Paternal condition: nABX vs CON
Activated
Suppressed
aerobic electron transport chain
mitochondrial ATP synthesis coupled electron transport
ATP synthesis coupled electron transport
proton motive force-driven ATP synthesis
mitochondrial respirasome
oxidative phosphorylation
inorganic cation transmembrane transporter activity
transmembrane signaling receptor activity
molecular transducer activity
signaling receptor activity
Gene ratio
Count
P(adj)
Paternal condition: LPHS vs CON
Activated
Suppressed
organellar ribosome
mitochondrial ribosome
structural constituent of ribosome
ribosomal subunit
large ribosomal subunit
ribosome
transcription regulator complex
Gene ratio
Count
P(adj)

**Supplementary Figure 1. The impact of specific environmental exposures on paternal physiology and F1 blastocysts.**
(A) Line plot showing growth curves of FVB males (F0 fathers) across CON, nABX and LPHS treatments. Error bars indicate standard deviation of mouse weights at each timepoint. Unpaired t-test; ns: p-value non-significant; *: p-value < 0.05. (B) Kaplan-Meier plot showing male mice's survival during the 6 weeks of indicated environmental exposure. (C) Testis to bodyweight ratio. (D) Representative images of males with open body cavity showing overt physiological responses after 6-7 weeks of indicated treatment. (E) Images of full (top) and empty (bottom) male ceca after 6-7 weeks of indicated treatment. (F) Representative images of F1 FVB blastocyst embryos generated through IVF. (F-G) Analysis of F1 blastocyst embryos with Copas Vision Sorter. (G) Scatter plot showing the time of flight (TOF) and Extinction of each blastocyst embryo colour by condition and by batch of experiments. Violin plot showing quantification of TOF or extinction. Unpaired t-test; ns: p-value non-significant. (H) Perimeter, Roughness and mean greyscale of F1 FVB embryos (see methods). Unpaired t-test; ns: p-value non-significant (I) Bar chart depicting efficiency of blastocyst development used for SMART-seq across multiple batches in the FVB background. (J) Table with numbers of embryos collected for SMART-seq (top) and passed quality thresholds for SMART-seq bioinformatic analysis (bottom). (K) Dot plot showing depth of blastocyst sequencing obtained across batches coloured by paternal condition. (L) Grouped bar chart showing number of male and female blastocysts across batches. (M) Upset plot showing overlap of DE genes between nABX-derived and LPHS-derived blastocysts. P-values are calculated using Fisher's exact test. (N) Bubble plot showing gene set enrichment analysis (GSEA) in nABX-derived (left) and LPHS-derived (right) blastocysts. + indicates activated pathways upon paternal treatment, - indicates suppressed pathways upon paternal treatment. (O) Principal Component Analysis (PCA) on genes identified to be DE in either nABX-derived or LPHS-derived blastocysts coloured by paternal treatment. Shaded histograms represent relative density of blastocysts along PC1 (top) and PC2 (right).

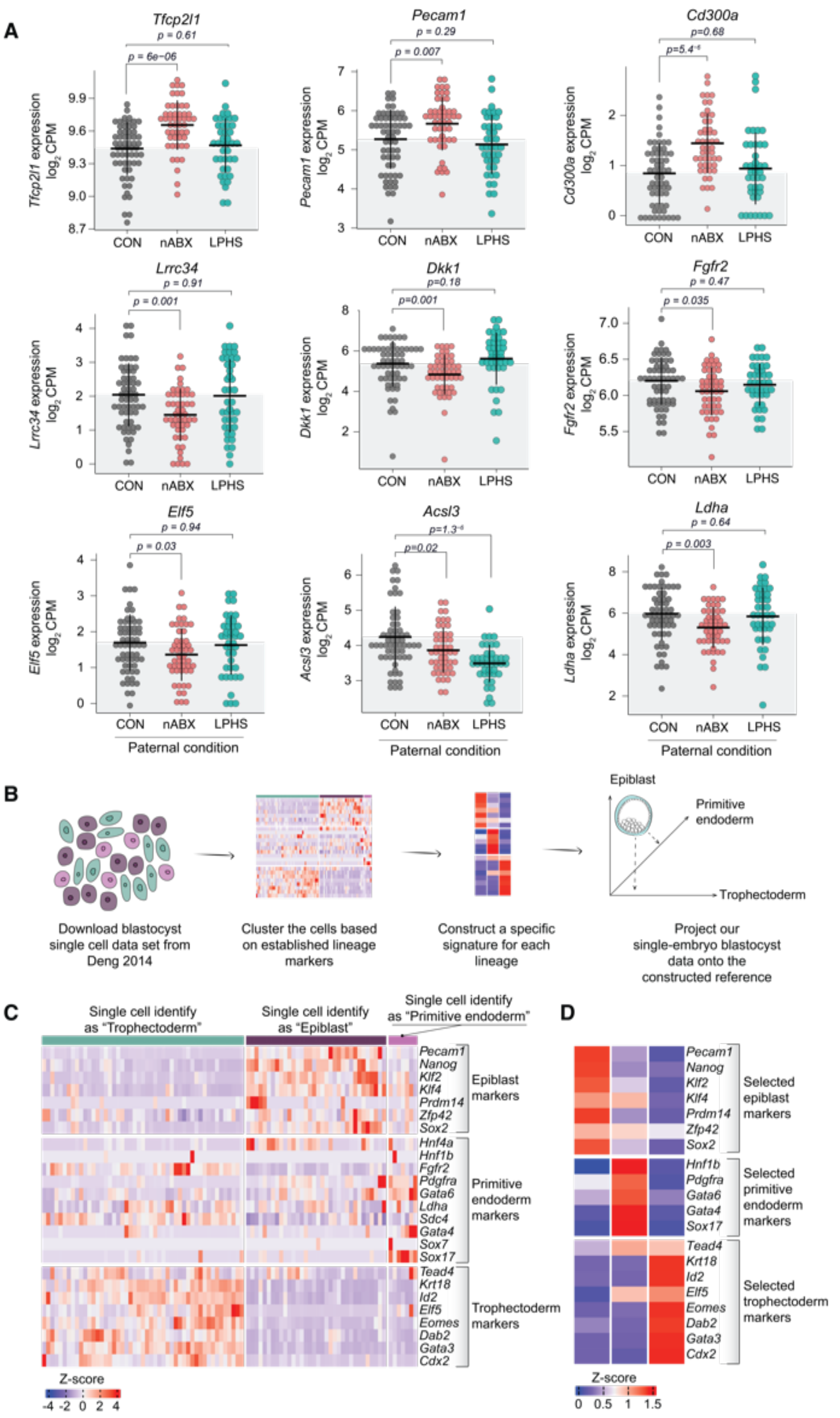


**Supplementary Figure 2. Characterising the IEI lineage-associated impact in F1 blastocysts.**
(A) Dot plots showing expression of lineage-associated DE genes in F1 blastocysts, driven by at least one paternal condition (from left to right and top to bottom): *Tfcp2l1*, *Pecam1*, *Cd300a*, *Lrrc34*, *Dkk1*, *Fgfr2*, *Elf5*, *Acsl3* and *Ldha*. *P-values* were computed using a two-tailed Wilcoxon test. (B) Schematic explaining the strategy to obtain lineage-specific signatures from published single-cell data (Deng, 2014) and projecting our blastocyst single-embryo dataset onto this reference (shown in Fig 2G-H). (C-D) Heatmaps of single-cell (C) and averaged (D) Z-scores of selected lineage-specific markers. Markers used to compute expression of lineage markers in embryos (Fig 2D-F and Fig 4C).

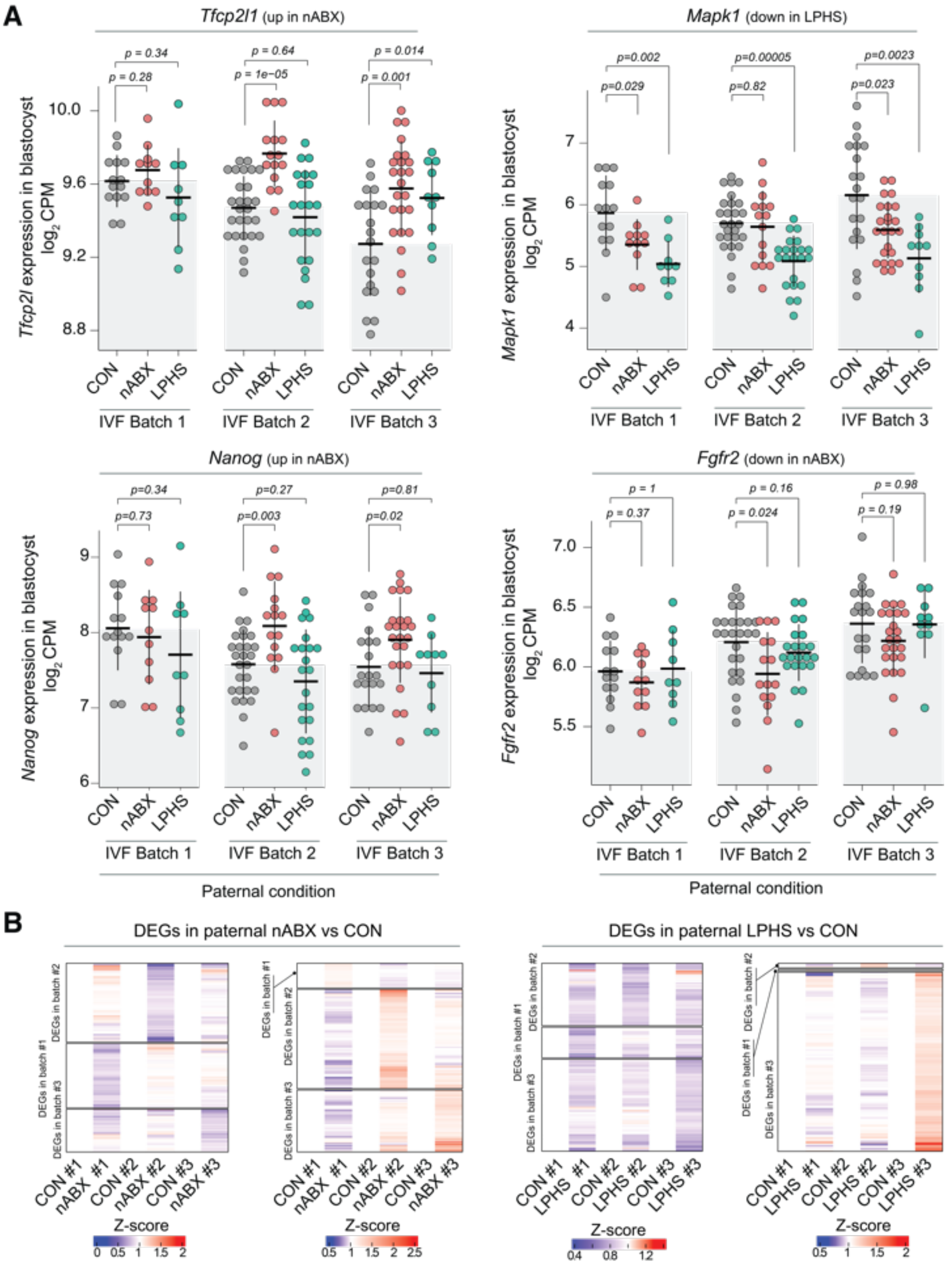


**Supplementary Figure 3. Assessing batch-effects in IVF-derived F1 blastocysts.**
(A) Dot plots showing log expression of genes in in F1 blastocysts derived from the indicated paternal condition, stratified by experimental batch: *Tfcp1l1, Nanog* (epiblast markers), *Mapk1 and Fgfr2 (signalling componants)*. Each datapoint indictaes a single blastocyst. *P-values* were computed using a two-tailed Wilcoxon test. (B) Heatmap showing fold changes of batch-specific DE genes relative to controls across all batches, split by batch of origin of the DE genes.

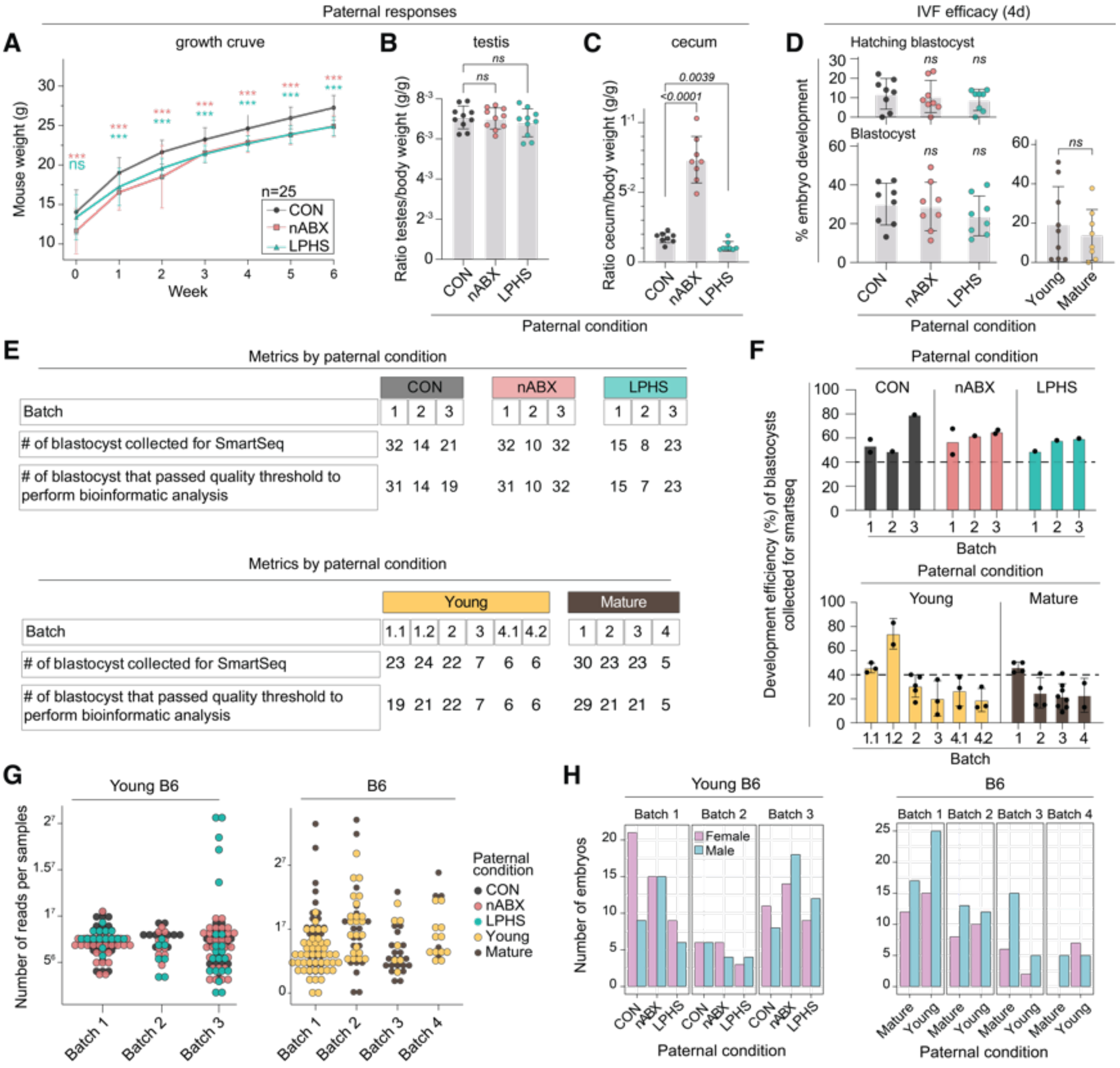


**Supplementary Figure 4. Interrogating the effect of age and genetic background on the IEI response in F1 blastocysts.** (A) Line plots showing growth curve of B6 males across CON, nABX and LPHS paternal treatments. Error bars indicate standard deviation at each timepoint. (B) Dot and bar plots showing testis / body weight ratio for B6 males after treatment. Error bars indicate standard deviation. (C) Dot and bar plots showing caecum / body weight ratio for young B6 males after treatment. Error bars indicate standard deviation. (D) Dot and bar plots showing rate of F1 blastocyst development and hatching from B6 across paternal treatment (left) and across ages (right). Error bars indicate standard deviation at each timepoint. (E) Tables showing numbers of embryos collected for SMART-seq and passed quality thresholds for SMART-seq bioinformatic analysis from B6 across paternal treatment (top) and across ages (bottom). All batches shown. (F) Bar charts showing developmental efficiency of blastocysts collected for SMART-seq across batches from B6 (top) and across ages (bottom). All batches are shown. Each dot represents the efficiency of blastocyst development in each culture well. Boxes represent the mean, and error bars indicate the standard deviation. (G) Dot plots showing depth of blastocyst sequencing obtained from young B6 fathers across paternal treatment (left), and across paternal ages (right). All batches are shown. (H) Grouped bar chart showing number of male and female blastocysts obtained from young B6 fathers across paternal treatment (left), and across paternal ages (right). All batches are shown. (A-D) P-values were computed using an unpaired t-test; [ns: not significant; *: p-value < 0.05; **: p-value < 0.005; ***: p-value < 0.0005].

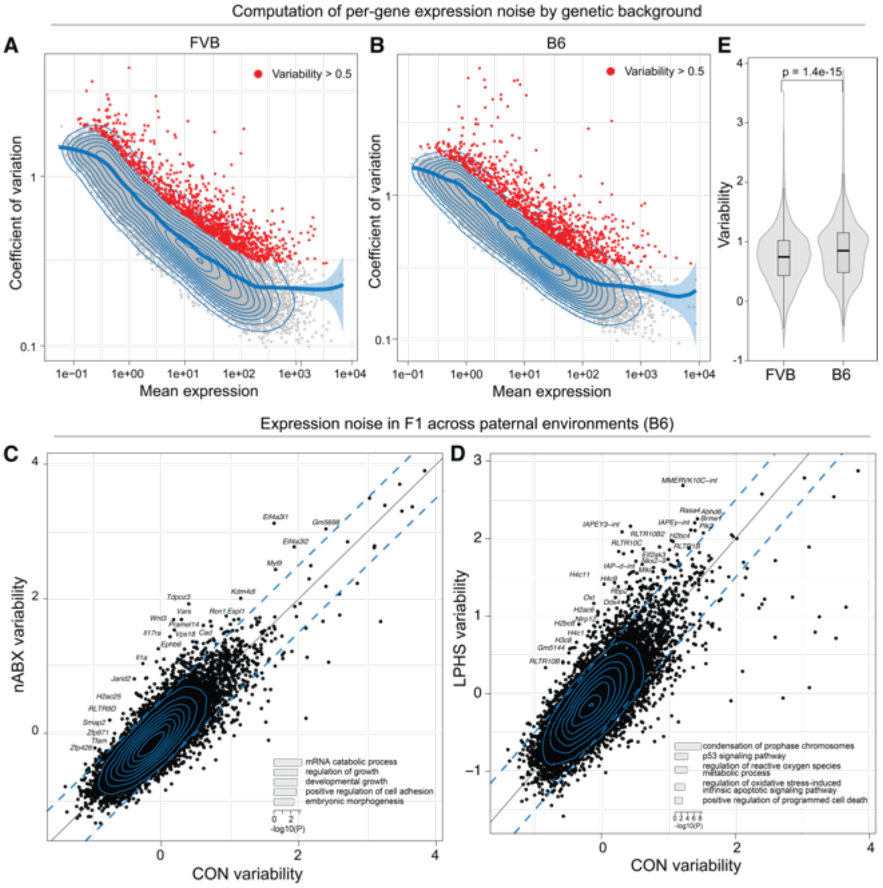


**Supplementary Figure 5. Gene expression variability across FVB and B6 backgrounds.**
(A-B) Scatter plots plotting coefficient of variation (CV) versus mean expression in FVB (A) and B6 (B) blastocysts. Blue line represents a generalised additive model (GAM) fit to the CV vs mean expression distribution. Red dots indicated genes with a variability score > 0.5 (see Methods). (C-D) (top) Scatter plots contrasting the variability of genes in the nABX-derived blastocysts (C) and LPHS-derived blastocysts (D) relative to controls in the B6 background. Genes showing higher variability in blastocysts of treated males are highlighted. (bottom) Horizontal bar chart highlighting enriched Gene Ontology (GO) terms in genes showing high variability in nABX-derived blastocysts (left) and LPHS-derived blastocysts (right) relative to controls in the B6 background. (E) Violin and box plot demonstrating the relative distribution of variability between the FVB and B6 backgrounds. P-value was computed using paired two-tailed T-test.

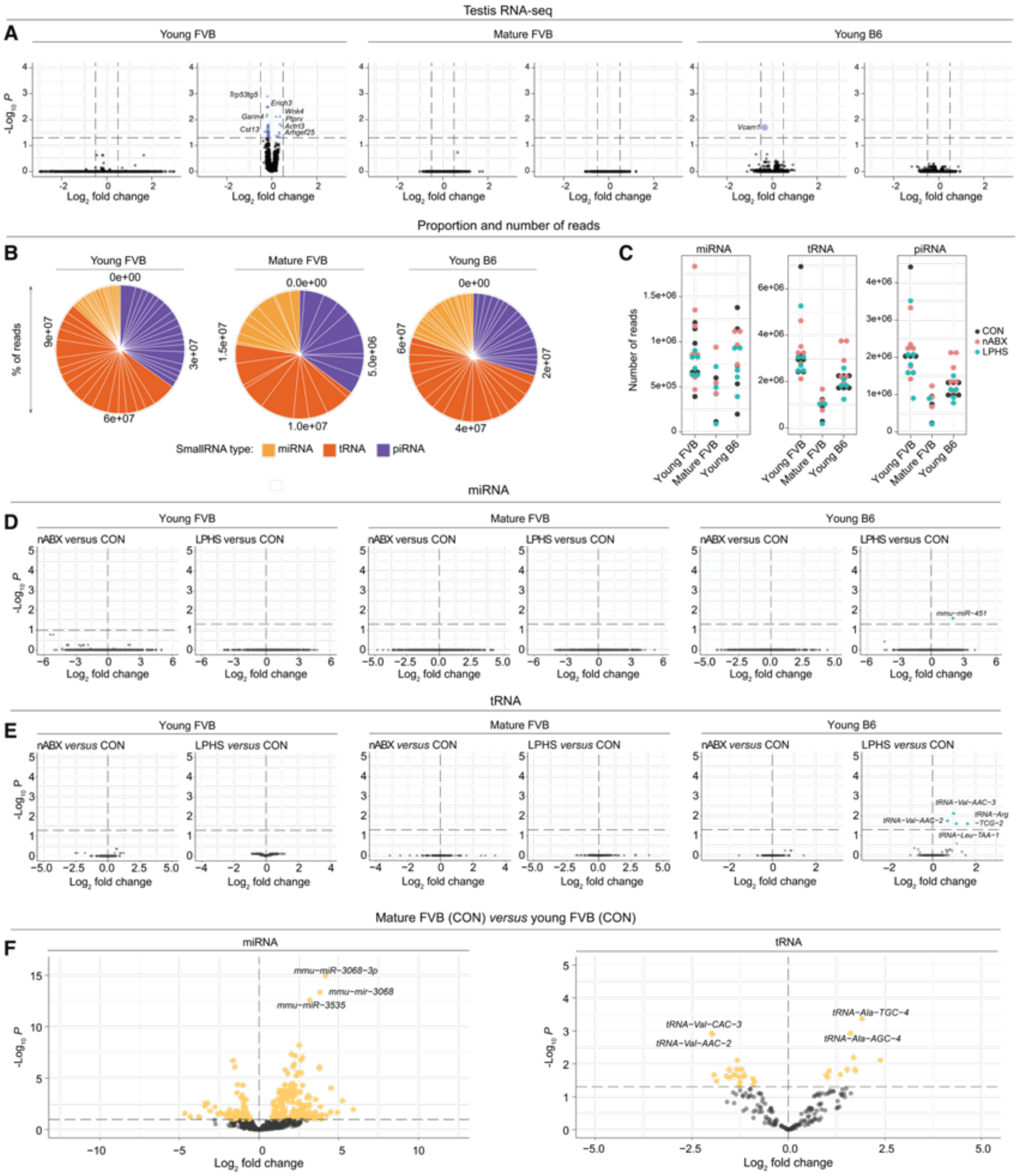


**Supplementary Figure 6. Paternal reproductive response to nABX, LPHS and aging across genetic backgrounds.**
(A) Volcano plots depicting differential gene expression in testis transcriptomes upon nABX and LPHS treatments in young FVB (left), mature FVB (middle) and young B6 (right) genetic backgrounds. (B) Pie charts showing relative distributions of different types of small RNAs in sperm of young FVB (left), mature FVB (middle) and young B6 (right) males. (C) Dot plots showing depth of small RNA sequencing split into miRNA (left), tRNA (middle) and piRNA (right) reads split across genetic background and coloured by condition. (D) Volcano plots of differential expression of miRNAs in sperm of nABX (left) and LPHS (right) males relative to controls in young FVB, mature FVB and young B6 genetic background. (E) Volcano plots of differential expression of tRNAs in sperm of nABX (left) and LPHS (right) males relative to controls in young FVB, mature FVB and young B6 genetic background. (F) Volcano plots of differential expression of miRNAs (left) and tRNAs (right) in sperm of untreated (CON) mature FVB versus young FVB males. (A, D-F) All volcano plots plot adjusted p-values versus $log_2$ fold-change values. Thresholds for significance are set at adjusted p-value < 0.05.
